# Synteny identifies reliable orthologs for phylogenomics and comparative genomics of the Brassicaceae

**DOI:** 10.1101/2022.09.07.506897

**Authors:** Nora Walden, M. Eric Schranz

**Affiliations:** Biosystematics Group, Wageningen University, Wageningen, The Netherlands; Centre for Organismal Studies, Heidelberg University, Heidelberg, Germany

**Keywords:** Ancestral genome, Angiosperm353, Brassicaceae, OrthoFinder, ortholog, paralog, phylogenomics, synteny

## Abstract

Large genomic datasets are becoming the new normal for any facet of phylogenetic research, but identification of true orthologous genes and exclusion of problematic paralogs is still challenging when applying commonly used sequencing methods such as target enrichment. Here, we compared conventional ortholog detection using OrthoFinder with ortholog detection through genomic synteny in a dataset of eleven representative diploid Brassicaceae whole genome sequences spanning the entire phylogenetic space. We then evaluated the resulting gene sets regarding gene number, functional annotation, gene and species tree resolution. Finally, we used the syntenic gene sets for comparative genomics and ancestral genome analysis. The use of synteny resulted in considerably more orthologs and also allowed us to reliably identify paralogs. Surprisingly, we did not detect notable differences between species trees reconstructed from syntenic orthologs compared other gene sets, including the Angiosperm353 set and a Brassicaceae specific target enrichment gene set. However, the synteny dataset comprised a multitude of gene functions, strongly suggesting that this method of marker selection for phylogenomics is suitable for studies that value downstream gene function analysis, gene interaction and network studies. Finally, we present the first ancestral genome reconstruction for the Core Brassicaceae predating the Brassicaceae lineage diversification ∼25 million years ago.

## Introduction

Sequencing data used for molecular phylogenetics has increased manifold over the last 30 years. Recent studies often use target enrichment, a cost-efficient method to simultaneously obtain nucleotide sequences of multiple genes from many taxa (Cronn et al. 2012; Weitemier et al. 2014). The bait sets for target enrichment may contain hundreds of genes and can be used for diverse sets of taxa, such as for a specific groups such as the Brassicaceae (Nikolov et al. 2019) or for larger groups such as for all angiosperms (Johnson et al. 2019). However, orthology, not paralogy of each gene (i.e., the homologous gene copy in each taxon is related by linear decent (Fitch 1970)) is required for many downstream analysis to obtain an accurate species tree, for example using ASTRAL (Zhang et al. 2018). Apart from obtaining an incorrect tree topology, inclusion of paralogs for species tree inference can also lead to erroneous older divergence time estimates (Siu-Ting et al. 2019; Zhou et al. 2022). Orthology is not trivial to achieve in target enrichment studies. Software such as orthoMCL (Li et al. 2003) or OrthoFinder (Emms & Kelly 2019) is often used to cluster genes from few available whole genome sequences or transcriptomes in the study species into groups of presumed orthologous genes (orthogroups) based on sequence similarity, and subsequently single-copy genes are selected for phylogenetic reconstruction (Baker et al. 2022; Johnson et al. 2019; One Thousand Plant Transcriptomes Initiative 2019). Orthology of the obtained sequences is then inferred from the nucleotide or amino acid sequence, ideally with additional filtering criteria (e.g., based on sequence identity and length) to exclude paralogs (Johnson et al. 2016). However, whether the orthogroups reliable only contain orthologs and no paralogs, and what the effects of mixed alignments on inferred species trees are, is rarely assessed.

Gene and genome duplications, gene loss, and variable rates of molecular evolution are all potential sources of error for ortholog detection. Error rates are highest for genes with high evolutionary rates, in particular for those with high between site rate heterogeneity, leading to clustering of many incomplete orthogroups (Natsidis et al. 2021). Single-copy gene families are thus predominantly comprised of genes with conserved nucleotide sequence and gene function (De Smet et al. 2013; Li et al. 2016) such as those involved in photosynthesis and cell cycle, while genes involved in local adaptation, such as those related to response to stimulus or signal transduction, are underrepresented (Li et al. 2017). This limits the subsequent use of phylogenomic target enrichment datasets. However, more and more research projects aim to investigate trait evolution in a phylogenomic context. For such studies, larger gene sets of reliable orthologs covering a wider range of gene functions would be highly advantageous, so that they can be used both to reconstruct the phylogenetic tree and subsequently identify the molecular basis of trait differences between taxa.

When whole genome sequences are available, additional information apart from nucleotide or amino acid sequence can be used to detect orthologous genes. Synteny, the collinearity of genes across genomes, could help identify reliable orthologs, because conserved gene order is expected between orthologous blocks. Additionally, many paralogs are derived from transposition to different locations in the genome. Notable exceptions are paralogous blocks that originated from ancient whole genome duplication (WGD), a phenomenon common in plants (e.g., Bowers et al. 2003; Vanneste et al. 2014; One Thousand Plant Transcriptomes Initiative 2019). The duplicated genes may subsequently undergo fractionation, through which one or the other gene copy is lost, or subfunctionalization, which may be linked to relaxed selection (Cheng et al. 2018). Both may hinder the correct identification of one-to-one orthologs, since a WGD-derived paralog may be wrongly added to a group of orthologous genes following loss of the respective ortholog, or clustering may add multiple paralogs from some taxa. However, syntenic paralog blocks derived from WGDs can easily be identified based on synonymous substitution rate (Ks).

The Brassicaceae are a relatively large angiosperm family with ∼ 4000 species in 350 genera (Walden, German, et al. 2020). With genome sequences available for the model plant *Arabidopsis thaliana* and many important crop species such as cabbages and rapeseed, the family has become a model system for genome evolution in recent years. However, the family’s phylogeny has so far proven difficult to resolve despite considerable efforts. The branching order of the main lineages differs between plastid based phylogenies (Walden, German, et al. 2020; Hendriks et al. 2022) and phylogenies based on nuclear genes (Nikolov et al. 2019; Huang et al. 2016; Kiefer et al. 2019; Hendriks et al. 2022), and support values for deeper nodes can be low. All Brassicaceae share the At-α WGD (Bowers et al. 2003; Schranz & Mitchell-Olds 2006) that occurred after divergence from the sister family Cleomaceae ∼ 40 million years ago (Edger et al. 2015), and many Brassicaceae tribes have undergone additional meso- and neopolyploidizations (Hohmann et al. 2015; Mandáková, Li, et al. 2017). The genome structure of the family has been studied in detail in many species of different tribes, and a system of 22 genomic blocks A-X has been established based on synteny of the genomes of *A. thaliana, Arabidopsis lyrata, Capsella rubella* and *Brassica rapa* (Schranz et al. 2006; Lysak et al. 2016). With many genome sequences now available, the Brassicaceae provide an ideal study system to investigate whether the use of synteny information for ortholog identification could be beneficial for future phylogenomics studies.

We selected eleven diploid Brassicaceae species with whole genome sequences available covering all major evolutionary lineages (Fig. 1). We compare orthologs identified through synteny with single-copy orthogroups from OrthoFinder regarding the total number of genes identified, bootstrap support, ASTRAL quartet scores and resulting species tree topology. Furthermore, we assess differences between both methods using subsets of available target enrichment gene sets for angiosperms (Johnson et al. 2019) and Brassicaceae (Nikolov et al. 2019). We evaluate the use of larger gene sets for studying trait evolution by comparing number of gene ontology (GO) terms and protein classes found in every gene set. Finally, we show that the usefulness of synteny does not end with ortholog detection for phylogenomics, and reconstruct ancestral genomes at nine crucial nodes in the evolutionary history of the Brassicaceae family.

**Figure 1.**
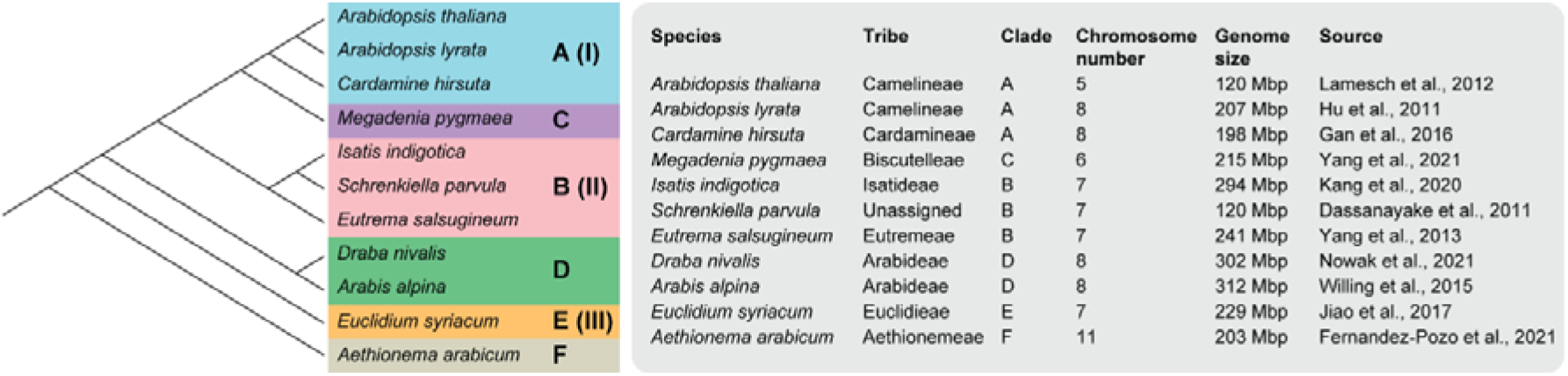
Brassicaceae species included in the study. The phylogenetic tree following Huang et al. (2016) with clades A-F (and lineage names I-III according to Walden et al. (2020)) is given on the left. Placement of *Megadenia* as sister to clade A followed Guo et al. (2021). Information on genome sequences for the studied species, including tribe and clade, chromosome number, genome size of the genome sequence and citation for the genome sequence are given in the right panel.

## Results

### Syntenic mapping identifies high numbers of orthologous genes

To reliably identify orthologs across Brassicaceae, we first used syntenic mapping of our selected 10 diploid genomes (Fig. 1) to *Arabidopsis thaliana*. We identified 21,221 genes with an ortholog in synteny in at least one of our ten other study species, and 7,825 genes with a syntenic paralog retained from the At-α WGD. Our further analyses were restricted to the 6,058 orthologs and 1,406 At-α paralogs (hereafter simply termed ‘paralogs’) that were found in synteny across all taxa, and we split them into five groups illustrated in Fig. 2a: syntenic orthologs without paralogs (‘no paralogs’), with syntenic paralogs retained only in some species (‘some paralogs’) or with syntenic paralogs in all species (‘syntenic paralogs’), and syntenic paralogs with syntenic orthologs retained only in some species (‘no syntenic orthologs’) or with syntenic orthologs across all species (‘syntenic orthologs’). As paralogs were only detected when orthologs were present there was no group containing paralogs without orthologs. Only 9.5% of orthologs (575 genes, 40.9% of paralogs) were also retained in synteny in paralogs in all species (Fig. 2b); an additional 1,650 orthologs had paralogs retained in synteny in some species (27.2% of orthologs); for 3,833 orthologs (63.3%) we did not detect a paralog in any species. Of the 1,406 syntenic paralogs, 831 (59.1%) did not have orthologous copies kept in synteny in all species.

**Figure 2.**
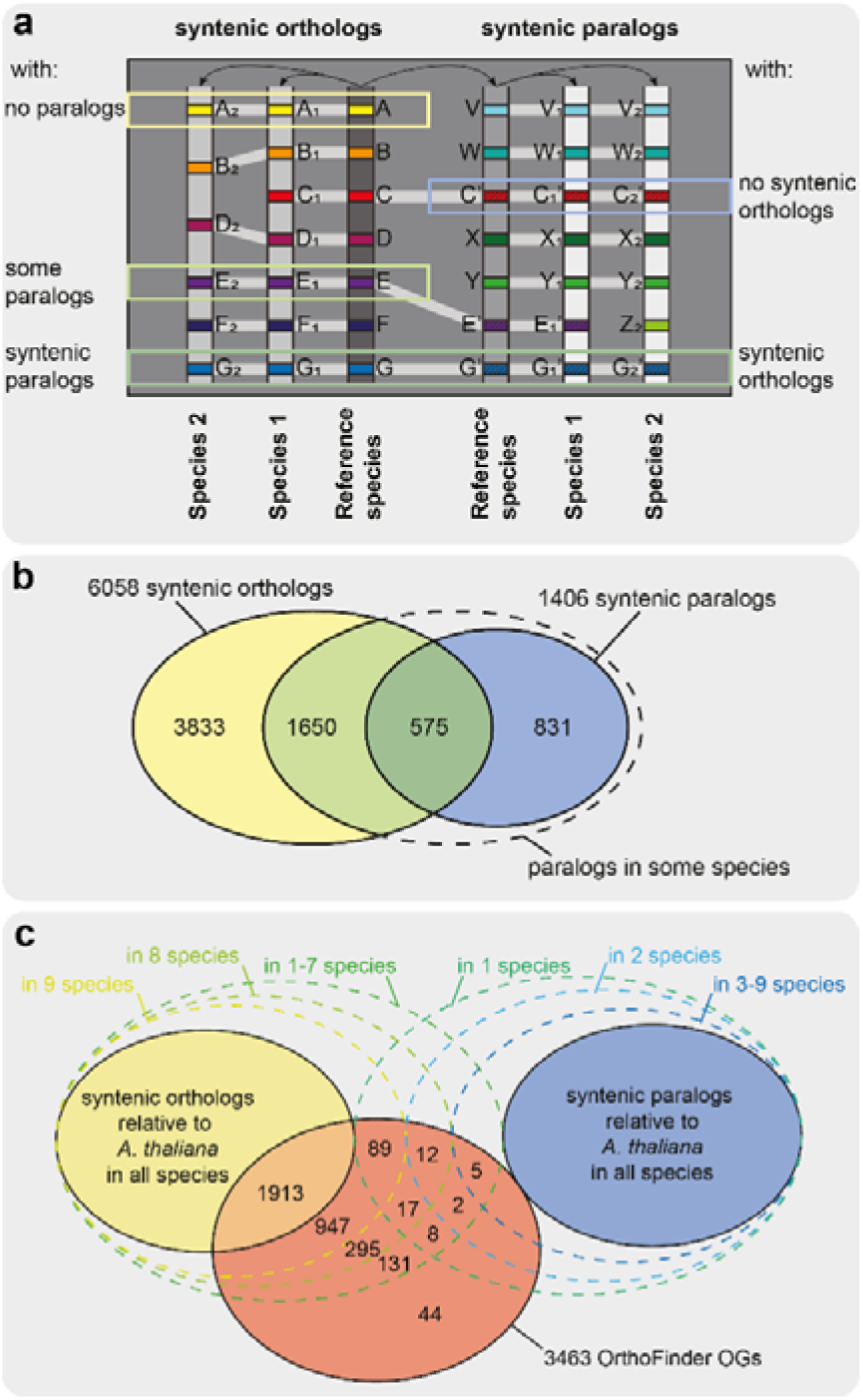
Distribution of syntenic orthologs/paralogs and OrthoFinder orthologs. a) Schematic drawing of syntenic mappings of two species against a reference. The following categories of orthologs/paralogs were analyzed: Syntenic orthologs across all species without syntenic paralogs (‘no paralogs’), with syntenic paralogs found in some species (‘some paralogs’), or with syntenic paralogs retained in all species (‘syntenic paralogs’); syntenic paralogs without syntenic orthologs detected in all species (‘no syntenic orthologs’) and syntenic paralogs with a complete set of syntenic orthologs (‘syntenic orthologs’). b) Number of syntenic orthologs/paralogs found across the eleven species of our study displayed as a Venn diagram. c) Venn diagram showing the number of single-copy OrthoFinder orthogroups found for each combination of orthologs/paralogs in synteny to A. thaliana and orthologs not in synteny. Few orthogroups had less than eight syntenic orthologs and are thus summarized in one category.

### Many OrthoFinder orthogroups are syntenic and few contain paralogs

For comparison, we used OrthoFinder to cluster homologous genes from all eleven species by sequence similarity. This resulted in 46,838 orthogroups, 3,463 of them strict single-copy orthogroups having a single gene copy in each taxon, which were the focus of our analysis. Using the information gained from syntenic ortholog and paralog identification, we assessed the composition of these single-copy orthogroups. As each orthogroup contains a single *A. thaliana* sequence, we searched for the genes from the other ten species among all syntenic orthologs and paralogs identified relative to *A. thaliana*. Genes that were not found may instead represent other paralogs such as transposed duplicates. No orthogroups were comprised solely of paralogs, but only 1,913 (55 %) had syntenic orthologs relative to *A. thaliana* in all ten species (Fig. 2c); this group represents the most reliable orthologs among OrthoFinder orthogroups. Most other orthogroups (1,242 or 35.9%) were comprised of syntenic orthologs in eight or nine species without any detected syntenic At-α paralogs, i.e., the other one or two genes in the orthogroup must be homologs located in a different genomic position. The two species not in synteny in the highest number of orthogroups were *Draba nivalis* and *Aethionema arabicum* (Fig. S1); generally, taxa more closely related to the reference species *A. thaliana* had more syntenic orthologs. Only 44 orthogroups (1.27%) contained no syntenic genes. The remaining 133 orthogroups (3.8%) were comprised of intermediate numbers of syntenic and other orthologs and paralogs, including paralogs retained from At-α but potentially also transposed duplicates or older paralogs.

We also investigated the overlap between our syntenic orthologs and paralogs with orthogroups from OrthoFinder, including multi-copy gene families. Most syntenic orthologs (5,807 or 95.9%) were contained within only a single orthogroup (Suppl. Fig. S2), indicating that across the ca. 30 million years of divergence among Brassicaceae, splitting orthologs into multiple orthogroups does not seem to be a widespread problem, at least when only considering genes with conserved genomic position. However, for syntenic paralogs, we detected 562 (40%) across multiple orthogroups, in line with the idea that sequence similarity-based clustering algorithms are more prone to errors for faster evolving gene copies such as paralogs, which may underlie less strict evolutionary pressure. In both groups, syntenic homologs were found across up to four different OrthoFinder orthogroups.

### Evaluation of gene sets for phylogenomics

To assess the impact of the two ortholog detection methods on inferred species trees, we first compared mean maximum-likelihood (ML) bootstrap support as a proxy for well-resolved gene trees. In addition to the five synteny gene sets from above, we also analyzed six of the larger sets of OrthoFinder orthogroups with eight, nine or ten syntenic orthologs and two, one or no paralogs. Mean bootstrap support among gene sets was in the range of 60.7-68.9%, with the highest value in the set of syntenic orthologs without paralogs, followed by OrthoFinder orthogroups containing only syntenic orthologs (Fig. 3a, b). The lowest mean bootstrap support values were found in the sets of OrthoFinder orthologs with only eight orthologs and one or two paralogs; however, sample size was small for these two sets. The subsets of syntenic and OrthoFinder orthogroups from the two target enrichment gene sets showed contrasting patterns: Gene trees from the Angiosperm353 set had relatively low mean bootstrap support (65.4 and 67.6%, respectively), while support was very high among gene trees from the Brassicaceae set (73.7 and 73.3%, respectively). The latter is likely due to implementation of stringent filtering criteria.

**Figure 3.**
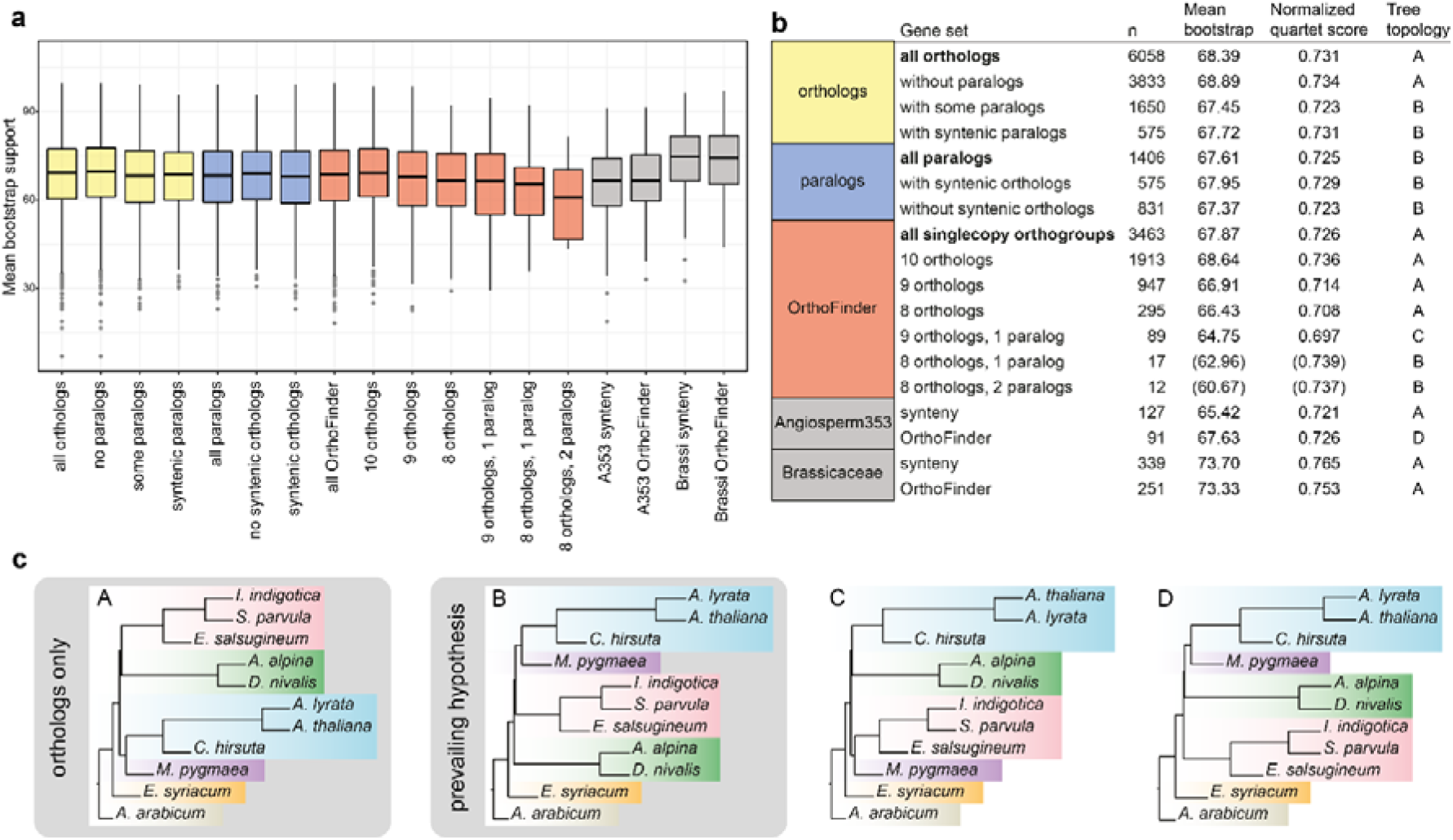
Evaluation of phylogenetic trees reconstructed from syntenic orthologs/paralogs and OrthoFinder orthogroups. a) Boxplot with mean bootstrap support for all trees of the respective gene sets. b) Information on gene set size, mean bootstrap support, normalized quartet score of the ASTRAL analysis against the species tree and the four different tree topologies (A-D) identified when running ASTRAL without constraint. c) The four tree topologies A-D.

**Figure 4.**
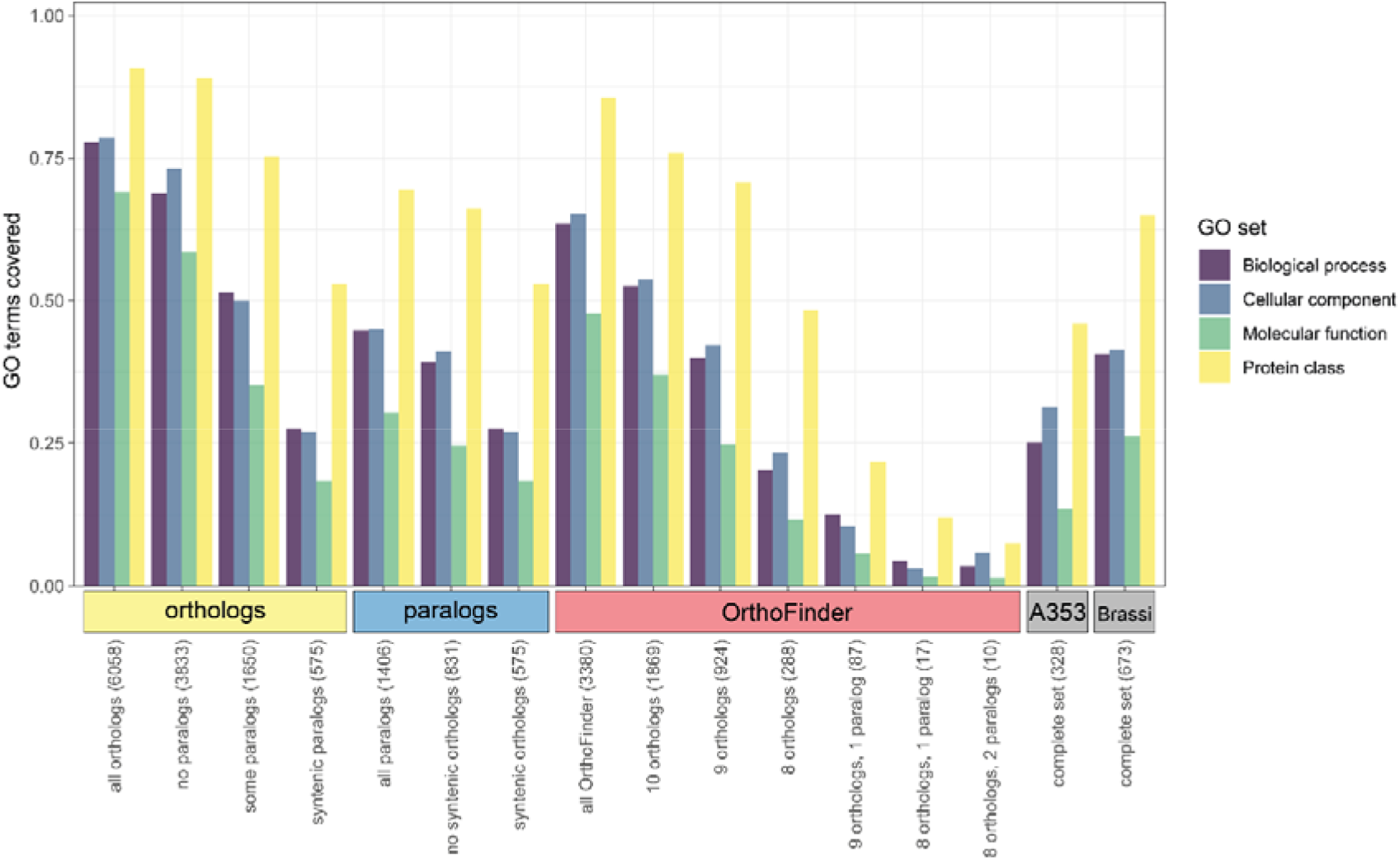
Fraction of GO terms and protein classes identified per gene set. GO terms and protein classes were identified using PANTHER, numbers are given relative to the total number of GO terms and protein classes relative to the A. thaliana annotation.

We also tested the effect of orthogroup selection on inferred species trees. First, we reconstructed species trees using ASTRAL (Zhang et al. 2018) for all gene sets separately. Gene sets with only orthologs showed a different branching order (topology A) from those that included paralogs (topology B, Fig. 3b, c). Interestingly, the syntenic ortholog subset of Angiosperm353 and Brassicaceae sets as well as the OrthoFinder subset of the latter also followed topology A, although the use of the entire Brassicaceae gene set in the respective study resulted in topology B (Nikolov et al. 2019), as did the combination of both sets (Hendriks et al. 2022). Additionally, we ran ASTRAL for all gene sets against the Brassicaceae phylogeny following the prevailing species tree hypothesis (also shown in Fig. 1) derived from three recent comprehensive phylogenomics studies (Huang et al. 2016; Nikolov et al. 2019; Hendriks et al. 2022) and compared the normalized quartet scores. The scores roughly followed the same trend as mean bootstrap support, with datasets comprised mainly of orthologs having higher values than those comprised of or including paralogs (with the exception of the two smallest gene sets, for which no reliable estimate could be obtained).

### Syntenic orthologs cover a large range of gene ontology (GO) terms

Next, we evaluated the coverage of GO terms and protein classes for each gene set relative to the complete annotation of the *A. thaliana* genome. The number of GO terms and protein classes in each gene set was related to gene number, with syntenic orthologs having the highest number of biological process, cellular component and molecular function related genes as well as the most different protein classes. When analyzing GO enrichment, we found that the fold enrichment was generally low (< 3-fold) for significantly enriched terms in larger gene sets, while small gene sets (e.g. Angiosperm353 or Brassicaceae) showed enrichment up to 50-fold, matching the expectation that the smaller gene sets comprise genes with a restricted subset of gene functions. The overrepresented terms also supported this idea; for example, plastid related cellular component terms were highly overrepresented in the Angiosperm353 set but less so in synteny-based gene sets (Suppl. Fig. S3). In line with previous studies on angiosperm core genes (Li et al. 2016), transcription factor related terms in the molecular function category as well as the protein class were underrepresented in both the Angiosperm353 and Brassicaceae set, while they were slightly overrepresented in most synteny- and OrthoFinder sets (Suppl. Fig. S4). Similarly, different terms related to stimulus response were highly underrepresented in the Angiosperm353 set in the biological process category (Suppl. Fig. S5). Interestingly, this set was also highly enriched for terms related to tRNA and rRNA processing. The larger gene sets however showed underrepresentation of defense related protein classes and the MADS-box transcription factors (Suppl. Fig. S6).

### Synteny data can serve as input for ancestral gene order reconstruction

Finally, we used the syntenic blocks as input for ancestral genome reconstruction. Our marker-based ancestral genome reconstruction used 336 markers containing blocks of syntenic genes present and in synteny in all eleven species across the Brassicaceae and consisted of 1-82 genes per marker, in total 3,392 genes. The average number of genes was 10 per marker, spanning 93 kb on average but ranging from 0.5 kb to 783.0 kb (Suppl. Fig. S7). As markers were required to be present in all species, the quality of the genome assemblies played a major role for marker coverage. For example, no synteny to *A. thaliana* could be detected for some sections of the *Euclidium syriacum* genome with the settings used here, and thus no markers were available for these parts of the genome (Suppl. Fig. S8). As a result, three of the shorter ABC blocks (G, P and T) are not covered in our reconstructions.

We reconstructed the ancestral marker order for all nine internal nodes (N1-N9) of the Brassicaceae phylogeny, using either *E. syriacum* or tribe Arabideae as the first diverging lineage (Suppl. Fig. S9) in the input phylogeny guiding the analysis, resulting in 18 reconstructions in total. As telomere position was inferred in the analysis, the number of chromosomes could be deduced from the number of telomeres present in each reconstruction. Between 12 and 17 telomeres were inferred, indicating chromosome numbers between seven and nine. Six to 15 CARs (Contiguous Ancestral Regions) were reconstructed by the algorithm, and where more CARs than inferred chromosomes were reconstructed, we combined them manually to obtain a likely version of the ancestral genome at the respective node (Fig. 5). Reconstructions at nodes N3-9 were identical for both input phylogenies. At five nodes (N3-7), the reconstruction only contained CARs with telomeres at both ends, indicating complete chromosomes; at two nodes (N8, N9), two CARs contained only one telomere and were combined. In contrast, different results were obtained using the two input phylogenies at nodes N1 and N2 (Suppl. Fig S10). At node N2, 15 telomeres were reconstructed for both analyses, with ten CARs for N2E and eleven for N2A. Two possible combinations of CARs were found for N2A, with the lower half of chromosome 7 being either the same as in N2E or the unmatched CAR from N2E. At node N1, 17 telomeres were inferred for N1E, while 15 were found for N1A; distinct differences between the two reconstructions included the number of B-U adjacencies found on CB1/7 and the combination of CB2/3 into a single chromosome in N1A.

**Figure 5.**
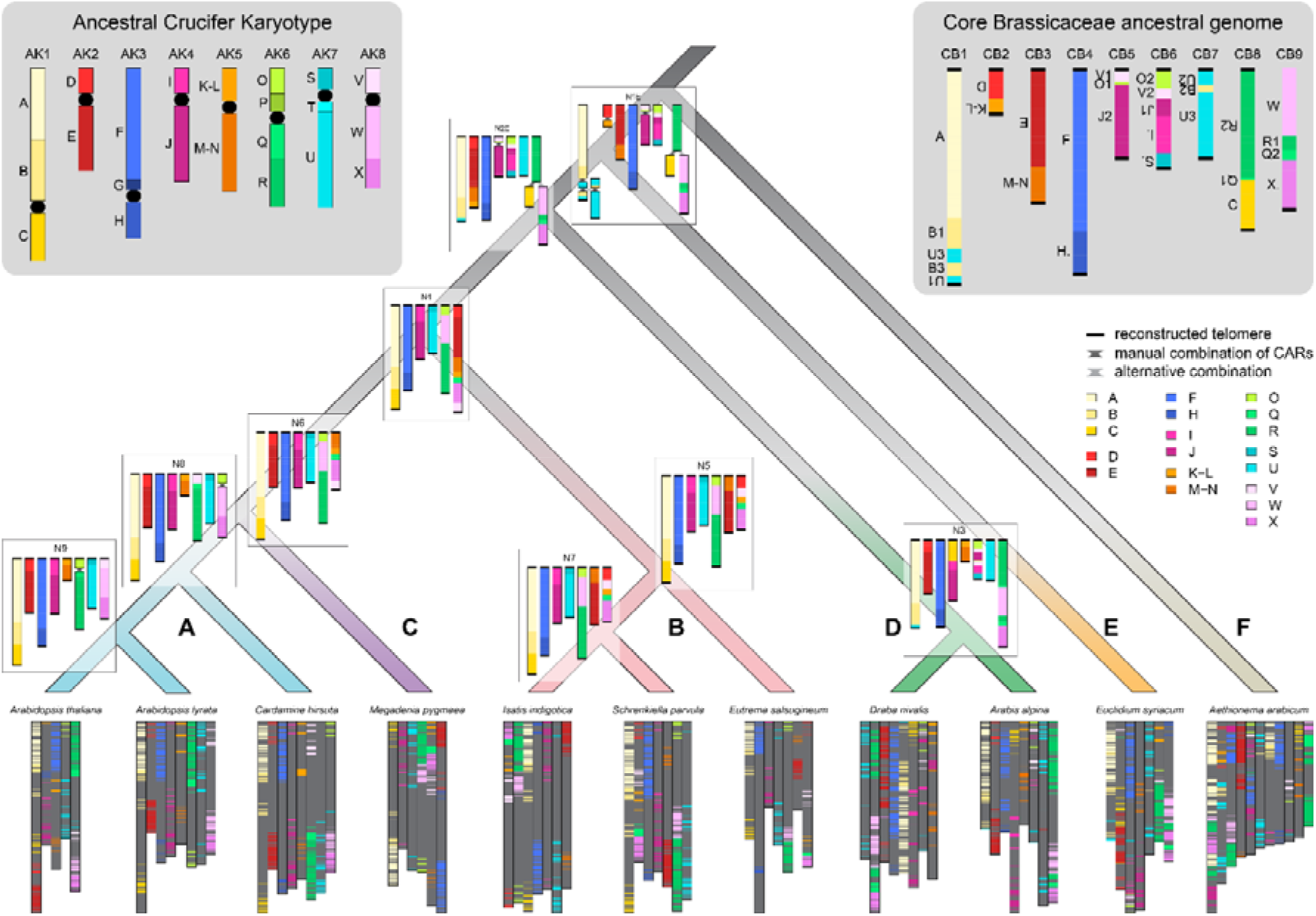
Schematic illustrations of ancestral genomes along the Brassicaceae phylogeny. Ancestral genome reconstruction was guided by an input phylogeny with *Euclidium syriacum* as the first diverging lineage (see Fig. 1). Markers are represented by horizontal bars colored by ABC blocks (Schranz et al., 2006; Lysak et al., 2016), black bars represent telomeres reconstructed by ANGES. CARs were combined manually for reconstructions containing CARs with less than two telomeres. Where more than one chromosome needed manual assembly, CARs were combined to retain marker adjacencies also found in related extant genomes; when multiple combinations were possible, the one found in *E. Syriacum* is displayed in dark grey, while the alternative combination is displayed in light grey. Note that the displayed length of chromosomes here is relative to marker number, not nucleotide length or gene number except for the extant genomes. The Ancestral Crucifer Karyotype (ACK; Schranz et al., 2006; Lysak et al., 2016) is shown in the top left corner for comparison; black circles represent centromeres. The Core Brassicaceae ancestral genome as reconstructed here is highlighted in the top right corner.

Ancestral genome reconstructions at more recent nodes closely resembled extant genomes from the respective clades: Nodes N3 (Arabideae, clade D), N5/7 (clade B) and N8/9 are all identical to extant genomes or showed at most a single rearrangement. Notably, the reconstructed ancestor of *Arabidopsis* (N9) was identical to the ACK (Fig. 5). Chromosomes AK1, AK3, AK4 and AK7 from the ACK were conserved throughout clades A, B and C with minor inversions only and are also found in the reconstructed genome of their most recent common ancestor. At the next deeper node however, only AK3 was conserved as a unit, while all others showed some rearrangement or fusion. Common rearrangements between chromosomes involved AK6/8, AK2/5 and AK1/7, and all of them can be seen in the Core Brassicaceae ancestral genome (Fig. 5). In this most recent common ancestor of all Brassicaceae but tribe Aethionemeae, seven of 22 ABC blocks are broken into two pieces, or four in the case of block U, and are found on different chromosomes while twelve blocks are intact relative to the ACK. To highlight the rearrangements relative to the ancestral genome of Core Brassicaceae we show genomic blocks colored by CAR in N1E in Suppl. Fig. S11.

## Discussion

Paralogs are homologous gene copies that originate from gene or genome duplication (Fitch 1970). For phylogenomic studies, the inclusion of paralogs in gene trees gives problems that hinder the reconstruction of accurate species trees. Gene and genome duplications often lead to neo- or subfunctionalization of gene function, fueled by the relaxed constraints on sequence mutation that can arise with existence of a second gene copy (Cheng et al. 2018; Birchler & Yang 2022). However, duplicated genes may subsequently be lost, and this phenomenon occurs even after millions of years (Johri et al. 2022). Phylogenetic trees reconstructed from genes where one taxon is represented by a paralog derived from gene duplication with subsequent loss of the ortholog may not follow the species tree topology and divergence times estimation from this gene may not be accurate either (Siu-Ting et al. 2019; Zhou et al. 2022). Most species tree reconstruction models thus require and assume orthology, and phylogenomic studies only select orthologs *a priori*.

Potential paralogs can be excluded at different stages of phylogenomic analysis. Many target enrichment studies begin by selecting only single- or low-copy genes during probe design (e.g. Johnson et al. 2019; Nikolov et al. 2019). However, strict exclusion of all genes with paralogs is rarely possible or limits the number of potential target genes considerably when the taxa of interest have undergone WGDs in their recent history, which is common for plants, and many genes have paralogs (Ufimov et al. 2022). Subsequent filtering for paralogs during analysis of the sequencing data is often conducted, for example in one of the most commonly used target enrichment analysis pipelines, HybPiper (Johnson et al. 2016), based on sequence length and similarity. Paralogs can also be detected based on phylogenetic trees (Kocot et al. 2013), and the problem of hidden paralogy through early gene duplication followed by late gene loss can be overcome to some extent by filtering gene trees for presence of known monophyletic clades (Siu-Ting et al. 2019). On the other hand, some more recent approaches explicitly use paralogs for species tree reconstruction. The inclusion of genes consisting of both orthologs and paralogs may result in accurate species tree reconstruction when methods accounting for incomplete lineage sorting (Yan et al. 2021) or gene duplications (Zhang et al. 2020) are used. Alternatively, detection of paralogs can be conducted based on patterns of sequence divergence between alleles and paralogs within a sample to obtain separate alignments for orthologs and paralogs from older WGDs, which may contain phylogenetic signal and thus contribute to overall better phylogenetic resolution at the species tree level by including more gene trees (Ufimov et al. 2022).

In recent years, synteny has emerged as a source of genomic information for comparative genomics (Conover et al. 2021; Lovell et al. 2022; Zhao et al. 2017; Zhao & Schranz 2017, 2019) and phylogenomics (Zhao et al. 2021). Here, we make use of positional information to identify reliable orthologs for species tree reconstruction and evaluate the resulting gene sets with respect to gene numbers, functional annotations, gene and species trees, as well as their further use for comparative genomics. One of the most important advantages for using synteny is that paralogs are easily identifiable. In the context of whole-genome sequences they can either be identified by position (transposed paralogs are located in a different genomic context) or by sequence divergence (paralogs retained from WGD have higher Ks). We thus found a large number of orthologs, representing 22.3 % of all genes annotated in *Arabidopsis thaliana*, and were also able to reliably identify many At-αparalogs as well as filter out other paralogs. Interestingly, both the Angiosperm353 and Brassicaceae set of target enrichment genes performed well regarding the exclusion of paralogs. This is likely due to strict downstream filtering, leading to the selection of highly conserved genes that are preferentially retained in single-copy and are also under strong selective constraints and thus have low substitution rates.

The genes in our gene sets encoded different protein classes, were involved in a wide variety of biological processes, had many molecular functions and were located in various cellular components. Enrichment analysis also showed generally low fold-enrichment (< 3-fold) of many GO terms, suggesting in turn a high coverage of many functional categories. Few categories were underrepresented in synteny sets; interestingly Type 1 MADS box transcription factors were among underrepresented protein classes for two sets of syntenic ortholog genes (‘all orthologs’, ‘orthologs without paralogs’), in line with a recent study showing that some MADS-domain transcription factors are frequently transposed in the Brassicaceae (Madrid et al. 2021). In contrast to the diverse gene functional annotations in our synteny gene sets, single-copy genes and thus gene sets selected for phylogenomics using conventional methods often have a conserved nucleotide sequence and gene function, such as photosynthesis and DNA metabolism (Li et al. 2017, 2016).

We compared our Brassicaceae synteny gene set with a previously published set of target enrichment genes designed for phylogenomic analysis of Brassicaceae (Nikolov et al. 2019). The Brassicaceae gene set contained 673 genes (2.5 % of *A. thaliana*), thus unsurprisingly a smaller fraction of functional categories was covered. Interestingly, different GO categories were enriched in the two sets, likely due to the strict filtering criteria used to exclude paralogs in the Brassicaceae set. Specifically, more conserved genes such as those involved in tRNA modification were overrepresented in the Brassicaceae set, and less conserved transcription factor related terms were underrepresented. This pattern was even more striking when comparing our synteny gene sets with the Angiosperm353 set, which was specifically designed to include only single-copy genes and avoid genes with potential paralogs, since accurate filtering for paralogs may not always be feasible for large-scale analyses with such a broad taxonomic focus. The Angiosperm353 set was highly enriched for genes localized to the plastid, in line with previous studies showing enrichment for organellar localization among single-copy genes (Han et al. 2014). We thus conclude that the use of larger gene sets and different gene selection criteria for phylogenomics may provide genomic information allowing for downstream identification of traits under selection from target enrichment data.

Our analysis of gene and species tree resolution and topology showed surprisingly few differences between gene sets. Bootstrap values were not significantly different between most sets, with the exception of the Brassicaceae set, which had considerably higher mean bootstrap values, likely due to strict filtering criteria, as well as higher quartet scores compared to other gene sets. Species tree topologies generally fell into two groups. Trees based solely on orthologs had clades A and C as sister to tribe Arabideae and clade B, while trees based on genes with paralogs had tribe Arabideae as sister to a clade containing clades A, B and C. The differences in topology are likely caused by generally low quartet scores at deeper nodes in the Brassicaceae phylogeny, where first and second topology can be almost equally common (Suppl. Table S1). Due to the high number of genes used to reconstruct the species tree these nodes can have high posterior probabilities despite similar quartet scores for main and alternate topologies. The selection of genes may nonetheless influence the topology. We found that while the Brassicaceae species tree based on the complete Brassicaceae set as used in the original publication supports Arabideae as sister to clades A, B and C (Nikolov et al. 2019), the tree based on the syntenic subset we analyzed here followed the same topology we detected for other gene sets without paralogs, with clades A and C as sister to Arabideae and clade B. Interestingly, a recent genus-level phylogeny of Brassicaceae also found different topologies at deep nodes depending on filtering level, with the species tree inferred from the combined Angiosperm353 and Brassicaceae gene sets supporting topology B and stricter filtering resulting in a species tree more similar to our topology A (Hendriks et al. 2022). Altogether, we conclude that the use of synteny for selecting genes for phylogenomics does not necessarily perform better than more established methods when it comes to the topology of the resulting species tree with regards to tree resolution. However, for taxa with difficult to resolve relationships the method may aid in obtaining large sets of reliable orthologs. It would be interesting to compare our results to another species group with difficult backbone phylogeny. Our approach was aimed at identifying a large number of reliable orthologs for species tree reconstruction through conserved gene position. As it relies on high quality genome assemblies, its use for now is limited to taxa where such data is available. However, new genome assemblies are becoming available rapidly, and future studies should consider including synteny in their selection of genes for phylogenomic reconstruction.

Finally, we made use of the synteny blocks identified for phylogenomic analyses and reconstructed ancestral genomes for the Brassicaceae. Previous ancestral genome reconstructions in the Brassicaceae were limited by the availability of high-quality genome sequences. As only genomes from the two largest clades, A and B, were available, reconstructions should not be considered ancestral to the entire family; however, genomes from the earlier diverging lineages have become available over the past few years, now allowing us to reconstruct ancestral genomes from deeper nodes of the family. The Ancestral Crucifer Karyotype ACK with *n* = 8 (Schranz et al. 2006; Lysak et al. 2016) was the first published ancestral Brassicaceae genome, reconstructed from the genome structures of *Arabidopsis thaliana, Arabidopsis lyrata, Capsella rubella* and *Brassica rapa*. A later ancestral genome reconstruction added *Schrenkiella parvula*, but the reconstructed genome still greatly resembled the ACK (Murat, Louis, et al. 2015). In our study, the ancestor of *Arabidopsis* strongly resembles the ACK, while the ancestor of clade A has an additional translocation. This difference is likely due to our inclusion of *Megadenia pygmaea* as sister to clade A, as well as further outgroups showing similar adjacencies. Due to the lack of a non-Brassicaceae outgroup, we cannot obtain the ancestral genome of all Brassicaceae including first diverging tribe Aethionemeae; however, for the first time we reconstruct the genome of the MRCA of Core Brassicaceae from ∼ 25 million years ago (Walden, German, et al. 2020). Our newly reconstructed ancestral genome of Core Brassicaceae had a haploid chromosome number of *n* = 9. Interestingly, the ancestral chromosome number for Brassicaceae was estimated to be *n* = 7 (Carta et al. 2020) in an angiosperm wide context. It should be noted that the estimated number may be strongly impacted by the studied taxa and outgroups. This is likely true for the ACK with its strong influence from clade A, where *n* = 8 is the base chromosome number in most tribes, as well as in our study, where the chromosome numbers of representatives from earlier diverging lineages ranged from *n* = 7 in *E. syriacum* to *n* = 11 in *A. arabicum*. Furthermore, telomeres here were reconstructed based on telomere position in our eleven extant species, not based on sequence data. The addition of other genomes may thus change the number of chromosomes in reconstruction. Despite the uncertainties that arise from the reconstruction method itself, evidence for the correctness of the results can be found in extant genomes. For example, ancestral adjacencies of blocks B-U (Core Brassicaceae chromosome CB1) and O-V-J (CB6) are also found within paralogous blocks retained from At-α (Walden, Nguyen, et al. 2020) and can thus even be dated back to before this WGD event. In the future, the availability of a (diploid) genome from sister family Cleomaceae would allow us to potentially reconstruct the ancestral genome of all Brassicaceae including first diverging lineage Aethionemeae and further advance our understanding of genome evolution in the family.

## Material and Methods

### Taxon sampling and genomic resources

We selected diploid species with available high-quality genome assemblies representing the major taxonomic lineages of Brassicaceae following recent nuclear phylogenies (Nikolov et al. 2019; Huang et al. 2016): *Aethionema arabicum* (Fernande □ Pozo et al. 2021) from first-diverging tribe Aethionemeae (clade F), which was used as an outgroup here, *Euclidium syriacum* (Jiao et al. 2017) from clade E, *Arabis alpina* (Willing et al. 2015) and *Draba nivalis* (Nowak et al. 2021) from clade D, *Eutrema salsugineum* (Yang et al. 2013), *Isatis indigotica* (Kang et al. 2020) and *Schrenkiella parvula* (Dassanayake et al. 2011) from clade B, *Megadenia pygmaea* from clade C (Yang et al. 2021), and *Cardamine hirsuta* (Gan et al. 2016), *Arabidopsis lyrata* (Hu et al. 2011) and *Arabidopsis thaliana* (Lamesch et al. 2012) from clade A. For the phylogenetic position of *M. pygmaea* from tribe Biscutelleae we followed Guo et al. (2021) and placed it as sister to clade A. More information on genomic resources can be found in Fig. 1.

### Assembly of pseudo-chromosomes

The published genome sequences of *E. syriacum, E. salsugineum* and *S. parvula* were not assembled at the chromosome level yet. For *S. parvula*, assignment of the largest scaffolds to seven pseudo-chromosomes was already available (Dassanayake et al. 2011), while for the other two species we first manually generated pseudo-chromosomes by combining the longest contigs from the published genome assemblies using evidence from chromosome painting data. We obtained pairwise syntenic blocks of both species with *A. thaliana* using SynMap in CoGe (https://genomevolution.org/coge/; Lyons et al. 2008; Nelson et al. 2018) with default settings and assigned them to ABC blocks. We then used information from chromosome painting (Mandáková, Hloušková, et al. 2017; Mandáková & Lysak 2008) to infer the position and orientation of scaffolds. Finally, the scaffolds were combined into pseudo-chromosomes (separated by 100 N’s) and new, matching annotation files were generated for downstream analyses for all three species. The newly generated assemblies are available at CoGe (https://genomevolution.org/coge/) under accessions id61751 (*E. syriacum*), id61750 (*E. salsugineum*) and id61748 (*S. parvula*).

### Synteny detection and grouping of orthologs and paralogs

Pairwise synteny between *A. thaliana* and the other ten species was detected using SynMap with default parameters; synonymous substitution rates (ks) were calculated. The resulting syntenic blocks were combined into blocks of genes present and in synteny in all species using R 4.0.5 (R Core Team 2022). Bad syntenic matches (ks > 5) and resulting short blocks (block length < 5) were removed. Blocks were then split into orthologs, paralogs derived from At-αWGD (‘paralogs’ from here on), and all other blocks. Only orthologs and paralogs mapping to the main chromosomes were retained for further analysis. Filtering for paralogs was performed by median block ks determined for each species manually (see Suppl. Fig. S12). For *A. arabicum* and *E. syriacum*, where the ks peaks for orthologs and paralogs overlap slightly, assignments were adjusted manually for blocks with intermediate ks using ks, gene content and duplication status. Syntenic orthologs and paralogs were then assigned to their *A. thaliana* counterpart, and gene groups containing all eleven species were selected. A subset of orthologous genes had an At-αderived paralog. We also ran OrthoFinder version 2.4.0 (Emms & Kelly 2019) using the longest transcripts of each gene for all species as input to independently obtain single-copy orthogroups. Using the gene names, we then counted the number of orthologs and paralogs identified in synteny analysis in OrthoFinder orthogroups.

We compared our synteny and OrthoFinder gene sets to two target enrichment gene sets: (1) The Angiosperm353 set (Johnson et al. 2019) is meanwhile widely used for phylogenomic studies within and across angiosperm families and contains 353 genes. (2) A Brassicaceae specific gene set (Nikolov et al. 2019) was recently developed to resolve the phylogeny of the family and comprises 673 genes. From each of these two gene sets we selected syntenic orthologs and single-copy OrthoFinder orthogroups for comparison with our other gene sets.

### Sequence alignment and phylogenetic inference

For each orthogroup from synteny analysis or OrthoFinder, we first aligned the coding sequences of all eleven species using MACSE version 2.05 (Ranwez et al. 2018) to obtain a codon-aware nucleotide alignment. The nucleotide sequence was then curated using Gblocks version 0.91b (Castresana 2000) using codon mode and discarding nonconserved blocks. Maximum likelihood (ML) trees were reconstructed using RAxML version 8.2.12 (Stamatakis 2014) with GTR+ Γ model of rate heterogeneity and rapid bootstrap inference with 1000 replicates followed by thorough ML search.

Species tree reconstruction was performed using ASTRAL version 5.7.8 (Zhang et al. 2018). To compare gene sets obtained through synteny and orthology detection, we analyzed syntenic orthologs, paralogs and OrthoFinder gene sets separately. We also split the sets further to study whether the presence of a paralogous gene copy in syntenic orthologs or a paralog in an OrthoFinder orthogroup influenced the reconstructed phylogeny. For all sets, we ran ASTRAL first without any constraints, and then tested the data versus the hypothesized species tree (Fig. 1) to obtain comparable normalized quartet scores.

### Analysis of gene function

Gene ontology (GO) and protein class enrichment analysis was conducted using PANTHER v. 16.0 (Mi et al. 2021, released 2020-12-01). Enrichment was tested for each synteny and OrthoFinder gene set and additionally for the entire Angiosperm353 and Brassicaceae gene set, including genes that were not found in synteny or in OrthoFinder orthogroups, using all *Arabidopsis thaliana* genes as background. Significance was tested using the binomial test with Bonferroni correction for multiple testing.

### Ancestral genome reconstruction

The ancestral gene order was reconstructed with ANGES (Jones et al. 2012), as this software has previously been successfully used to reconstruct the ancestral genomes of eukaryotes, both animals (Neafsey et al. 2015) and plants (Murat et al. 2014; Murat, Zhang, et al. 2015). In short, the method rearranges user-specified orthologous markers such as genes or larger genomic regions found across species into CARs (Contiguous Ancestral Regions) and is guided by a bifurcating phylogenetic tree. As small rearrangements (e.g. transpositions) are common across many plant genomes and also among our selected species, the use of orthologous genes as markers does not result in long, contiguous output at chromosome or chromosome-arm level. Instead, we thus used longer syntenic blocks of genes present in all species. Markers were generated using pairwise synteny with *A. thaliana*. Syntenic maps were created using SynMap with parameters optimized for the following analysis steps. Default parameters were used except for the following: DAGChainer maximum distance between two matches was set to 10, DAGChainer minimum number of aligned pairs was set to 20, and synonymous substitution rates (ks) were calculated. The resulting syntenic blocks were combined into blocks of genes present and in synteny in all species using R v4.0.5 (R Core Team 2022). Only orthologous blocks mapping to the main chromosomes were considered. Filtering for paralogs and block optimization was performed as above. This resulted in 336 blocks containing 3,392 genes found in the same block in all species. The genomic coordinates of the blocks in each species were then used as input for ancestral genome reconstruction. To assess the extent to which the input phylogeny influenced the reconstructed ancestral genomes, we used two different input trees representing different hypotheses for the nuclear phylogeny: The representative of the first diverging lineage after tribe Aethionemeae was either *E. syriacum* (clade E, see Fig. 1) or the clade containing *A. alpine* and *D. nivalis* (clade D). The latter was chosen due to the proposed phylogenetic position of Arabideae or clade D as first diverging lineage based on similarities between the genome structures of *Arabis alpina* and *Aethionema arabicum* (Walden, Nguyen, et al. 2020). Markers were set as unique and universal, doubled markers were utilized to infer marker direction, and telomeres were added after greedy heuristic C1P optimization. For each of the two phylogenies, we reconstructed the ancestral genomes at all nine nodes, resulting in a total of 18 reconstructions.

### Combination of CARs to ancestral genomes

Ancestral genomes were manually assembled from the CARs obtained from ANGES following three rules: (1) Chromosomes end with telomeres. A CAR with reconstructed telomeres at both ends was thus considered a complete chromosome, while a CAR with one telomere was located at the chromosome end and could be combined with another one-telomere CAR to obtain a complete chromosome. (2) If a genome contained more than one chromosome made from two CARs, marker adjacencies in closely related extant lineages were considered, and CARs were combined in such a way that at least one extant adjacency was present in the ancestral reconstruction. (3) In cases where combination of CARs was ambiguous, we followed adjacencies in the respective first diverging lineage; however, alternative adjacencies are shown in Suppl. Fig. S10. For details, see Supplementary Methods.

The system of ABC blocks (Schranz et al. 2006; Lysak et al. 2016) that has been used to visualize Brassicaceae genomes for the past 15 years is based solely on the genomes of clades A and B. Here, we reconstructed the ancestral genome of Core Brassicaceae, the ancestor all major lineages except tribe Aethionemeae (clade F). To display the chromosomal rearrangements relative to this ancestor, we assigned a new coloring scheme based on the CARs and chromosomes of the ancestral genome at node N1E (Suppl. Fig. S11).

## Acknowledgements

This work was supported by the German Research Foundation [HO 6443/1 to N.W.].

## Author contributions

N.W and E.M.S designed the study, N.W. performed the analysis with input from E.M.S.,

N.W. drafted the manuscript with input from E.M.S.

## Notes

### Competing Interest Statement

The authors have declared no competing interest.

