## Supporting Information for "Synteny identifies reliable orthologs for phylogenomics and comparative genomics of the Brassicaceae"

#### Supplementary Methods

*Assembly of CARs into chromosomes*

In the following paragraphs, we describe in detail how we combined CARs from ancestral gene order reconstructions with ANGES into chromosomes for both input phylogenies. We generally followed three rules: 1) Chromosomes end in telomeres. A CAR with reconstructed telomeres at both ends represents a complete chromosome, while a CAR with one telomere is located at the chromosome end and a CAR with no telomere is generally located in the chromosome center. 2) If a reconstructed ancestral genome contains more than one chromosome made from two CARs, marker adjacencies in closely related extant species are considered. In those cases, we combined CARs in such a way that at least one extant adjacency is retained from the ancestral genome. 3) In cases where combination of CARs was ambiguous, we followed adjacencies in the respective first diverging lineage; however, alternative adjacencies are shown in Suppl. Fig. S10.

At five of nine reconstructions, all CARs were reconstructed with telomeres at both ends by the algorithm, requiring no manual assembly. At two nodes, only two CARs were reconstructed with a single telomere, which we combined into a chromosome. Finally, at the two most basal nodes, five or more CARs were reconstructed without telomeres at both ends. We combined CARs with at least one adjacency present in Arabideae, *Euclidium syriacum* or *Aethionema arabicum*. In all four cases, at least one alternative pairing of CARs was possible and is displayed in Fig. S10.

It should be noted here that our manual reconstruction of ancestral genomes is particularly prone to error (see Discussion) and should be regarded as merely one possible solution.

Node 9: ancestor of *Arabidopsis thaliana* and *Arabidopsis lyrata* (clade A)

At this node, results were identical for both input phylogenetic trees. ANGES reconstructed 16 telomeres, indicating eight chromosomes. This is in line with the expectations for the node, as the base chromosome number for most tribes in Lineage I/A is eight (Hohmann et al., 2015). The markers were assembled into nine CARs. Seven CARs were reconstructed with telomeres on each side and were thus considered full chromosomes; and we combined the remaining two CARs so that the telomeres were located at the chromosome ends. The genome structure strongly resembles that of the Ancestral Crucifer Karyotype ACK (Schranz et al., 2006; Lysak et al., 2016).

Node 8: ancestor of *Arabidopsis* and *Cardamine hirsuta* (clade A)

Also at this node, identical results were obtained for both phylogenetic trees. The number of reconstructed telomeres was 16 again, in line with base chromosome number for the lineage (see node 9). Seven of the nine CARs had telomeres at both ends and thus represented entire chromosomes, and the two CARs with only one telomere were combined into a chromosome manually. In general, the reconstruction was similar to the ancestor of *Arabidopsis*, with the exception of the reciprocal translocation between chromosomes 6 and 8 seen also in *Cardamine hirsuta*.

Node 7: ancestor of *Isatis indigotica* and *Schrenkiella parvula* (clade B)

Results were identical for analyses with both phylogenetic trees. Seven CARs were assembled with 14 telomeres, indicating seven chromosomes. This is in line with base chromosome number in many tribes of lineage II/B (Hohmann et al., 2015). The genome structure resembles that reported for tribes Isatideae, Sisymbrieae and Eutremeae previously (Mandáková and Lysak, 2008), with four chromosomes identical to the ACK and three chromosomes with large-scale rearrangements.

Node 6: ancestor of *Arabidopsis*, *Cardamine hirsuta* and *Megadenia pygmaea* (clades A+C)

Also at this node, the results obtained using the different phylogenetic trees as input were identical. Seven CARs were assembled with 14 telomeres, indicating seven chromosomes. *M. pygmaea* has six chromosomes, with D/E and F/H fused into one chromosome, O/W/R identical to that of the PCK and M-N/V/X/Q/K-L fused into another. The reconstruction shows a similar structure, but without the chromosomal fusion, and with a slightly different order of blocks on chromosome seven, where M-N and K-L are adjacent like in the ACK.

Node 5: ancestor of *Isatis indigotica*, *Schrenkiella parvula* and *Eutrema salsugineum* (clade B)

The reconstructed ancestral gene order for the three representatives of clade B was identical for both input phylogenies, and the same as the ancestor of *I. indigotica* and *S. parvula* (node 7).

Node 4: ancestor of species from clades A/B/C

At this node, the results were again identical for analyses with both phylogenetic trees. Six CARs with 12 telomeres were assembled, indicating six chromosomes. The reconstructed genome structure strongly resembles that of node 6 (clades A+B), but with ACK 2/3 fused.

Node 3: ancestor of *Draba nivalis* and *Arabis alpina* (clade D)

Also at this node, identical results were obtained for analyses with both phylogenetic trees. Eight CARs with 16 telomeres were reconstructed, indicating eight chromosomes. The reconstructed genome more closely resembles that of *Arabis alpina*, except for chromosome seven, which is more similar to *Draba nivalis*.

Node 2A: ancestor of clades A, B, C and E

At node 2, the two phylogenetic trees resulted in slightly different reconstructed ancestral genomes. With Arabideae (clade D) as first diverging lineage, eleven CARs were reconstructed, with a total of 15 telomeres, indicating more than seven chromosomes. Five CARs were reconstructed with telomeres at both ends and we thus consider them complete chromosomes. Five CARs had telomeres on one side, and one CAR consisting only of a single marker contained no telomeres. We manually assembled the remaining chromosomes so that every combination of CARs was supported by at least one adjacency in the extant genomes from clade D, E or F. For the small CAR with V-P and the CAR with J, only the combination of both was supported by an adjacency in extant genomes (in *Aethionema arabicum*). The combination of the single marker CAR containing a fragment of block Q with R was also found in the extant genomes of *A. arabicum* and Arabideae. For the remaining three CARs, two possible solutions were found: One, with block C as a separate chromosome and the CAR containing X-Q-R-W connected to Q-R as seen in extant genomes of Arabideae, the other one with X-Q-R-W as a separate chromosome and block C connected to Q-R, a connection seen in the extant genome of *Euclidium syriacum*.

Node 2E: ancestor of clades A, B, C and D

With *Euclidium syriacum* (clade E) as first diverging lineage, ten CARs were reconstructed, also with a total of 15 telomeres, indicating more than seven chromosomes. Again, five CARs were reconstructed with telomeres at both ends and considered complete chromosomes. The other five CARs each contained one telomere, and combination of CARs resulted in an ancestral genome reconstruction identical to that for Arabideae as first diverging lineage.

Node 1A: ancestor of Core Brassicaceae (clades A-E)

At node 1, the two phylogenetic trees resulted in different reconstructed ancestral genomes. With Arabideae (clade D) as first diverging lineage, eleven CARs with 15 telomeres were reconstructed. Five CARs contained telomeres on both ends and were considered complete chromosomes, five CARs contained only one telomere, and one CAR had no telomeres. While for the long part of block J the combination with O-V like in *Aethionema arabicum* was the only possible combination, three combinations were possible for R-Q: Either with V-O-J, resembling one of the adjacencies from *E. syriacum*, or with either of the CARs also combined with R-Q in N2 (W-R-Q-X as in Arabideae or C as in *E. syriacum*). Two combinations were also possible for C, the other one being with I-J-V-O, with the adjacency shared with Arabideae.

Node 1E: ancestor of Core Brassicaceae (clades A-E)

With *Euclidium syriacum* (clade E) as first diverging lineage, 15 CARs with 17 telomeres were found, indicating more than eight chromosomes. Three CARs contained two telomeres and were considered complete chromosomes. Eleven CARs contained one telomere and one CAR contained no telomere. A single supported adjacency was found for two combinations of CARs: D was combined with K-L like in *E. syriacum* and V-O was combined with J like in *A. arabicum*. R-Q was combined with either C or W-R-Q-X like in node 2.

### Supplementary Tables

**Table S1. ASTRAL quartet scores.** Scores are given for each node in the species tree reconstructed from the respective gene sets given on the left.

| | **Gene set** | ***Arabidopsis*** | **A** | **D** | **E&F** | **A&D** | **A&C** | **D&B** | **AC&B** | **AC&D** | **B&C** | **B** | ***Sparvula/ lindigotica*** |
|---|---|---|---|---|---|---|---|---|---|---|---|---|---|
| **synteny** | **all orthologs** | 0.98/0.01/0.01 | 0.82/0.09/0.09 | 0.98/0.01/0.01 | 0.47/0.31/0.22 | | 0.47/0.23/0.29 | 0.37/0.35/0.27 | | | | 0.85/0.07/0.08 | 0.85/0.07/0.08 |
| | no paralogs | 0.98/0.01/0.01 | 0.83/0.08/0.09 | 0.98/0.01/0.01 | 0.48/0.31/0.22 | | 0.47/0.23/0.29 | 0.38/0.35/0.27 | | | | 0.85/0.07/0.07 | 0.86/0.08/0.07 |
| | some paralogs | 0.98/0.01/0.01 | 0.8/0.1/0.1 | 0.97/0.02/0.01 | 0.46/0.3/0.23 | | 0.47/0.24/0.29 | 0.36/0.36/0.28 | | | | 0.84/0.08/0.08 | 0.84/0.07/0.09 |
| | syntenic paralogs | 0.98/0.01/0.01 | 0.83/0.09/0.08 | 0.96/0.02/0.02 | 0.45/0.31/0.25 | | 0.49/0.34/0.17 | | 0.36/0.36/0.28 | | | 0.86/0.08/0.06 | 0.86/0.07/0.07 |
| | **all paralogs** | 0.98/0.01/0.01 | 0.8/0.1/0.1 | 0.96/0.02/0.02 | 0.45/0.29/0.26 | | 0.47/0.37/0.16 | | 0.37/0.35/0.28 | | | 0.86/0.07/0.07 | 0.84/0.08/0.08 |
| | syntenic orthologs | 0.98/0.01/0.01 | 0.8/0.1/0.09 | 0.96/0.02/0.02 | 0.44/0.3/0.26 | | 0.48/0.36/0.16 | | 0.37/0.36/0.27 | | | 0.86/0.07/0.06 | 0.85/0.08/0.07 |
| | no syntenic orthologs | 0.98/0.01/0.01 | 0.8/0.1/0.1 | 0.96/0.01/0.02 | 0.45/0.27/0.28 | | 0.46/0.16/0.37 | | 0.36/0.35/0.29 | | | 0.85/0.07/0.07 | 0.84/0.08/0.09 |
| **OrthoFinder** | **all singlecopy orthogroups** | 0.99/0.01/0.01 | 0.81/0.09/0.1 | 0.98/0.01/0.01 | 0.47/0.31/0.22 | | 0.46/0.3/0.24 | 0.37/0.35/0.27 | | | | 0.84/0.08/0.08 | 0.85/0.08/0.07 |
| | 10 orthologs | 0.99/0/0.01 | 0.82/0.09/0.09 | 0.99/0/0.01 | 0.47/0.3/0.23 | | 0.47/0.31/0.22 | 0.37/0.36/0.27 | | | | 0.87/0.06/0.06 | 0.86/0.07/0.07 |
| | 9 orthologs | 0.98/0.01/0.01 | 0.81/0.08/0.1 | 0.98/0.01/0.01 | 0.47/0.31/0.23 | | 0.44/0.3/0.26 | 0.38/0.35/0.27 | | | | 0.8/0.11/0.09 | 0.85/0.07/0.09 |
| | 8 orthologs | 1/0/0 | 0.75/0.12/0.12 | 0.97/0.02/0.01 | 0.45/0.32/0.23 | | 0.45/0.28/0.27 | 0.38/0.32/0.3 | | | | 0.82/0.09/0.09 | 0.83/0.1/0.07 |
| | 9 orthologs, 1 paralog | 1/0/0 | 0.79/0.11/0.11 | 0.98/0.01/0.01 | 0.46/0.29/0.25 | 0.36/0.33/0.3 | | | | | 0.42/0.27/0.31 | 0.73/0.18/0.09 | 0.89/0.06/0.05 |
| | 8 orthologs, 1 paralog | 0.99/0.01/0.01 | 0.84/0.13/0.03 | 1/0/0 | 0.45/0.29/0.26 | | 0.49/0.18/0.33 | | 0.42/0.21/0.37 | | | 0.78/0.18/0.04 | 0.88/0.06/0.06 |
| | 8 orthologs, 2 paralogs | 1/0/0 | 0.79/0.17/0.05 | 0.99/0/0.01 | 0.44/0.29/0.26 | | 0.6/0.1/0.29 | | 0.51/0.36/0.13 | | | 0.6/0.23/0.17 | 0.92/0/0.08 |
| **Angiosperm353** | synteny | 0.99/0.01/0 | 0.87/0.06/0.07 | 0.98/0.01/0.01 | 0.46/0.22/0.32 | | 0.49/0.29/0.22 | 0.39/0.31/0.3 | | | | 0.79/0.09/0.12 | 0.86/0.08/0.07 |
| | OrthoFinder | 0.96/0.01/0.03 | 0.87/0.05/0.09 | 0.99/0/0.01 | 0.49/0.23/0.28 | | 0.46/0.35/0.19 | | | 0.36/0.35/0.3 | | 0.86/0.09/0.06 | 0.9/0.02/0.08 |
| **Brassicaceae** | synteny | 1/0/0 | 0.89/0.05/0.05 | 1/0/0 | 0.49/0.31/0.19 | | 0.54/0.26/0.2 | 0.39/0.36/0.25 | | | | 0.89/0.04/0.07 | 0.88/0.07/0.06 |
| | OrthoFinder | 1/0/0 | 0.88/0.06/0.05 | 1/0/0 | 0.49/0.21/0.3 | | 0.49/0.31/0.2 | 0.38/0.27/0.35 | | | | 0.88/0.07/0.05 | 0.88/0.08/0.04 |

#### Supplementary Figures

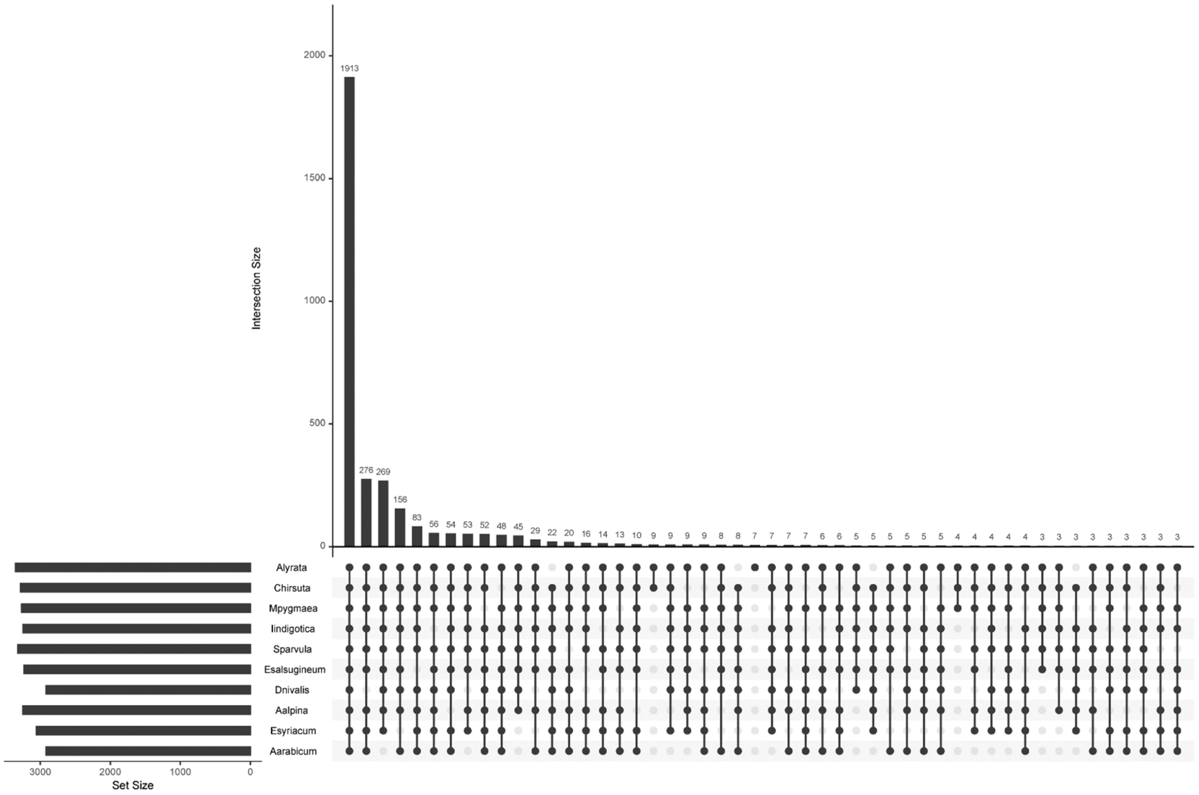


**Figure S1. Upset plot visualizing taxon content of OrthoFinder orthogroups.** The intersection size (y-axis) shows the number of orthogroups with each taxon set (x-axis); only the largest 50 sets are shown. The plot was generated using the R package UpSetR (Gehlenborg, 2019).

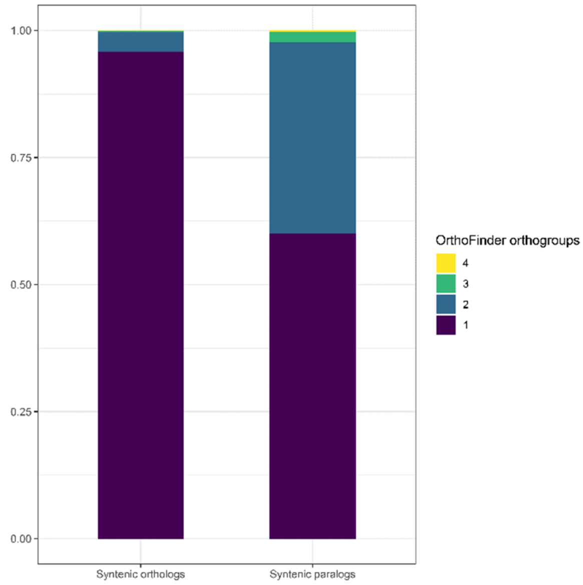


**Figure S2. Number of OrthoFinder orthogroups per syntenic set.** For each syntenic ortholog and paralog we checked across how many different OrthoFinder orthogroups the sequences from our eleven study taxa were found. Relative numbers are given.

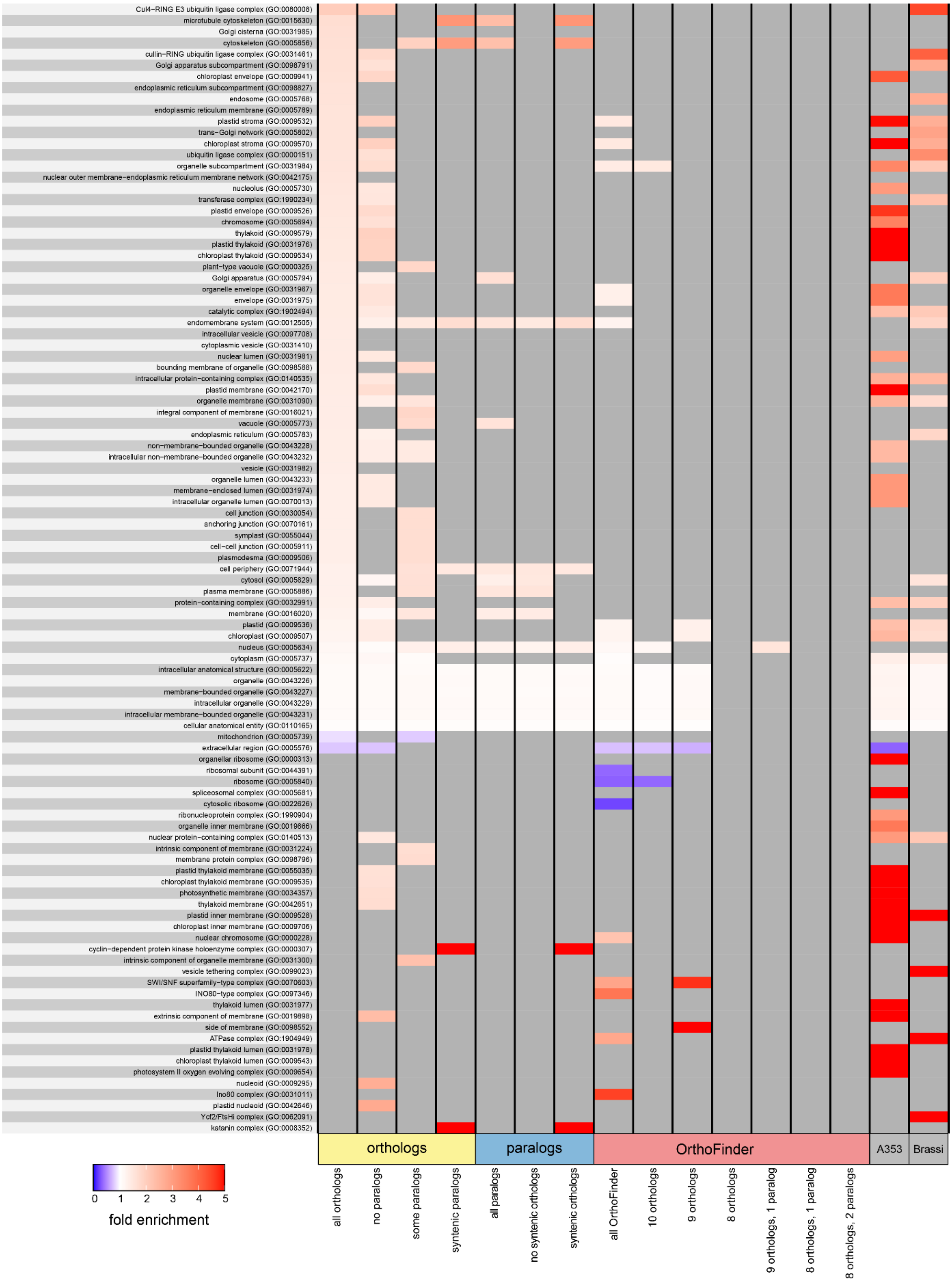


**Figure S3. GO enrichment for cellular component.** Fold enrichment for terms with significant enrichment (binomial test with Bonferroni correction for multiple testing; p-value < 0.05) are shown for all 16 gene sets. Heatmap was plotted using R package superheat v1.0.0 (Barter and Yu, 2022), fold enrichment > 5 was rare and thus merged.

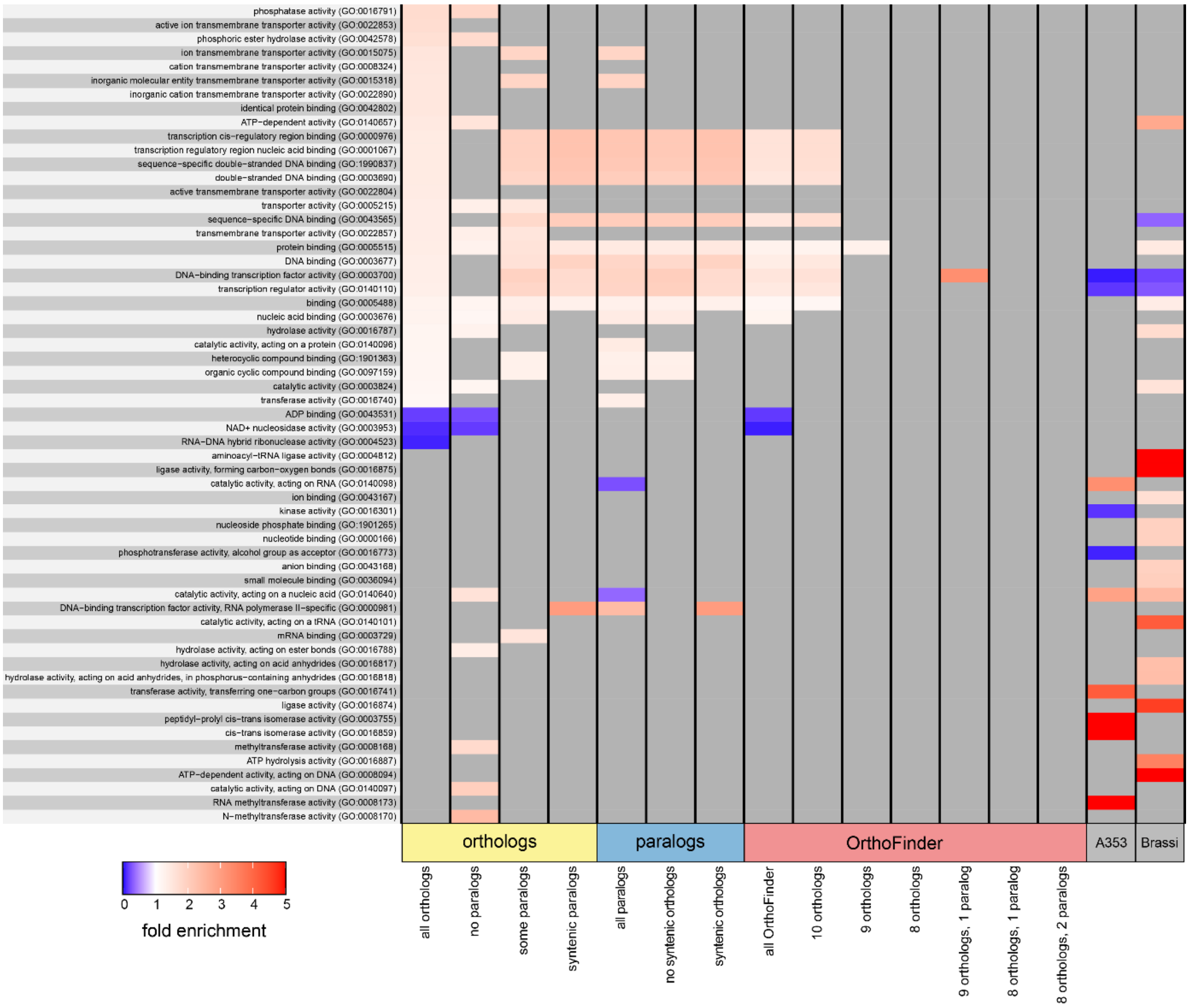


**Figure S4. GO enrichment for molecular function.** Fold enrichment for terms with significant enrichment (binomial test with Bonferroni correction for multiple testing; p-value < 0.05) are shown for all 16 gene sets. Heatmap was plotted using R package superheat v1.0.0 (Barter and Yu, 2022), fold enrichment > 5 was rare and thus merged.

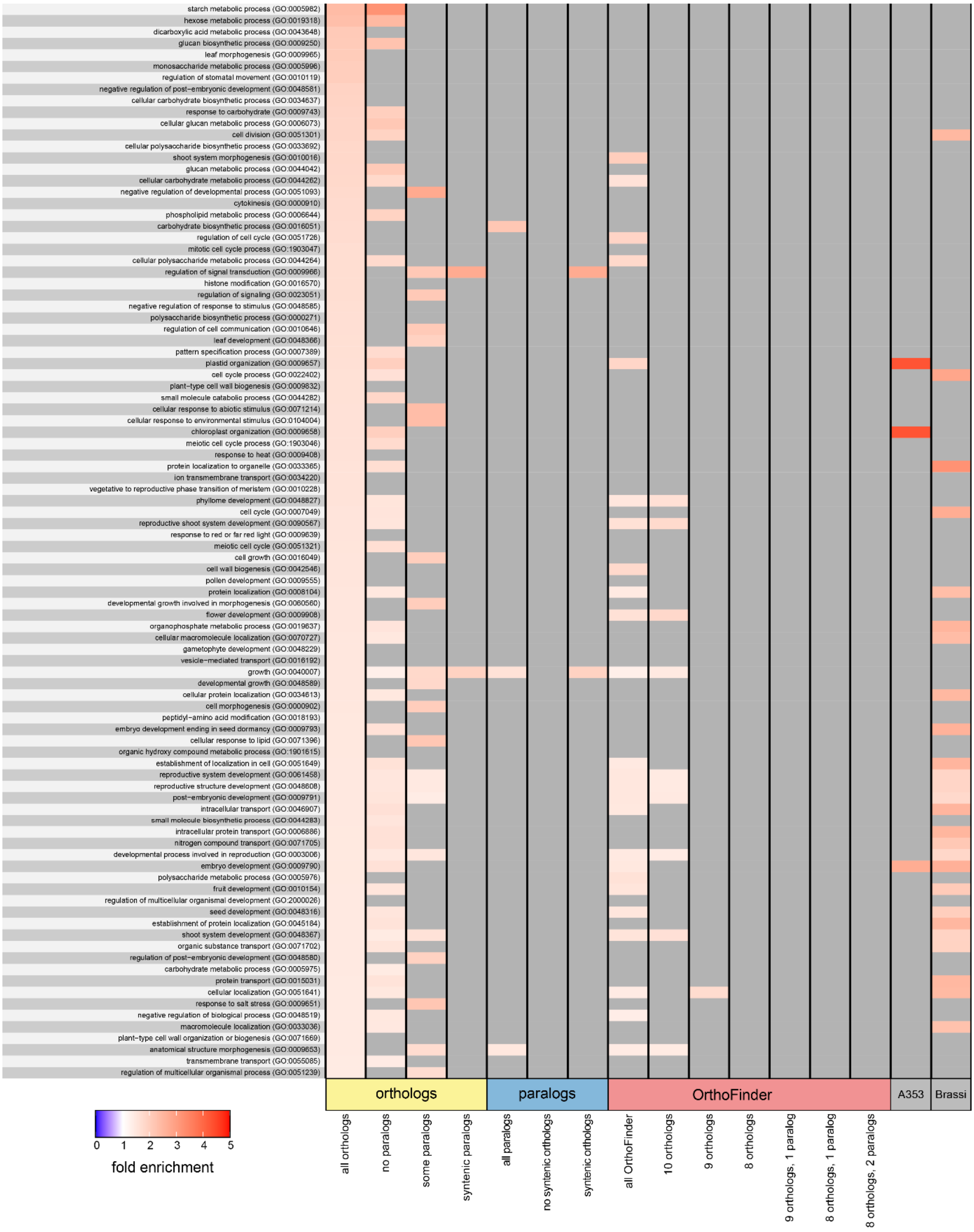


**Figure S5. GO enrichment for biological process.** Fold enrichment for terms with significant enrichment (binomial test with Bonferroni correction for multiple testing; p-value < 0.05) are shown for all 16 gene sets. Heatmap was plotted using R package superheat v1.0.0 (Barter and Yu, 2022), fold enrichment > 5 was rare and thus merged.

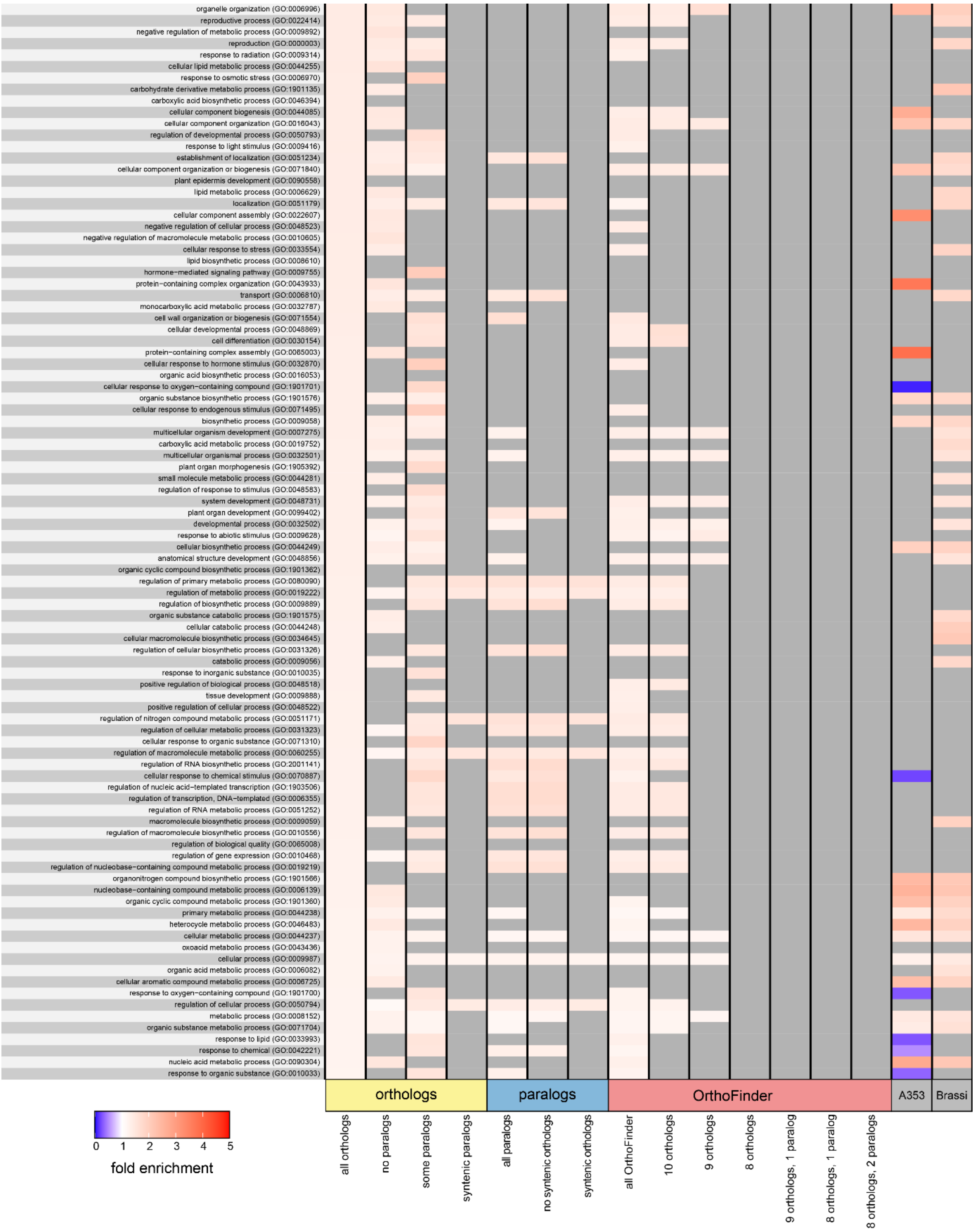


**Figure S5 (continued). GO enrichment for biological process.** Fold enrichment for terms with significant enrichment (binomial test with Bonferroni correction for multiple testing; p-value < 0.05) are shown for all 16 gene sets. Heatmap was plotted using R package superheat v1.0.0 (Barter and Yu, 2022), fold enrichment > 5 was rare and thus merged.

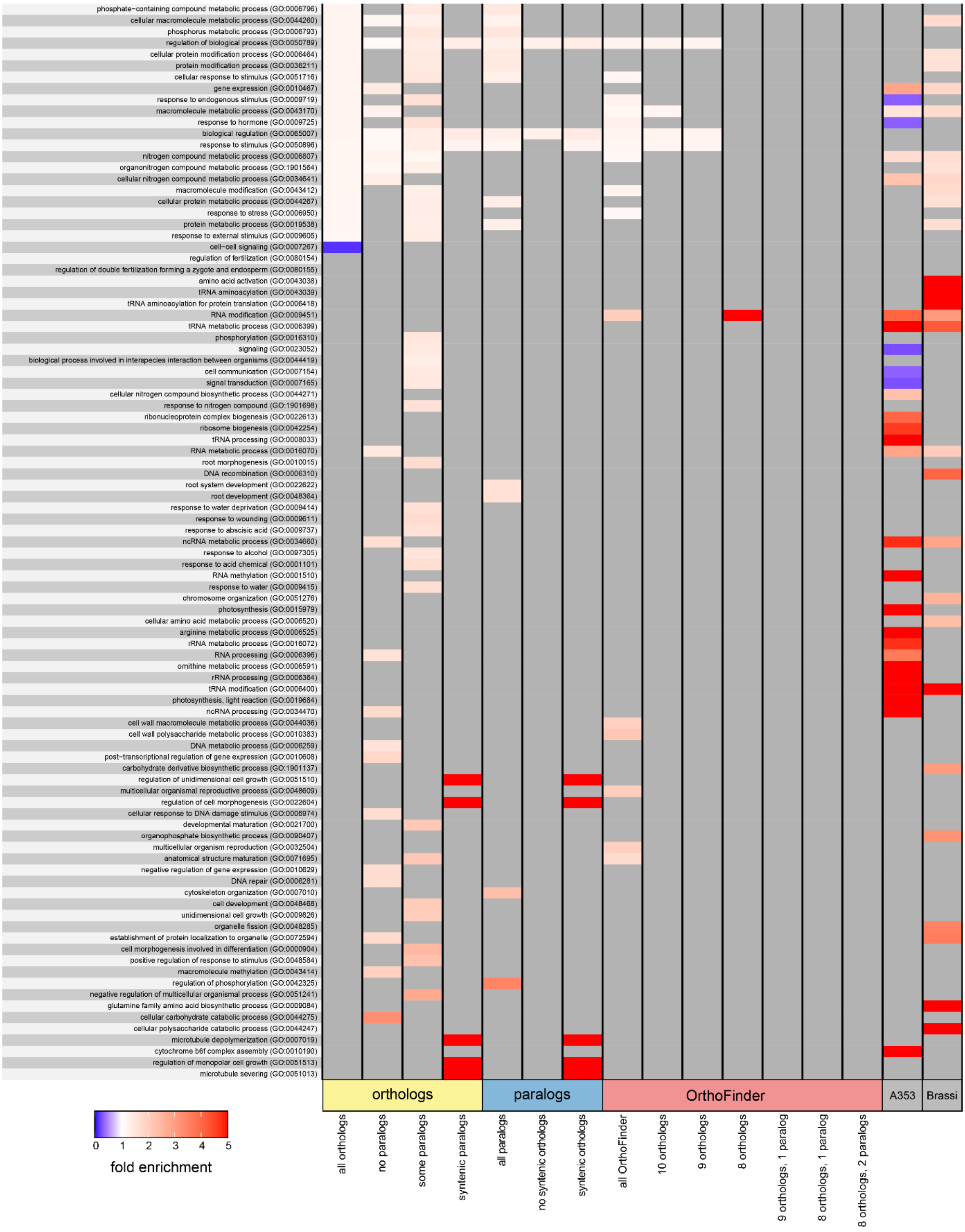


**Figure S5 (continued). GO enrichment for biological process.** Fold enrichment for terms with significant enrichment (binomial test with Bonferroni correction for multiple testing; p-value < 0.05) are shown for all 16 gene sets. Heatmap was plotted using R package superheat v1.0.0 (Barter and Yu, 2022), fold enrichment > 5 was rare and thus merged.

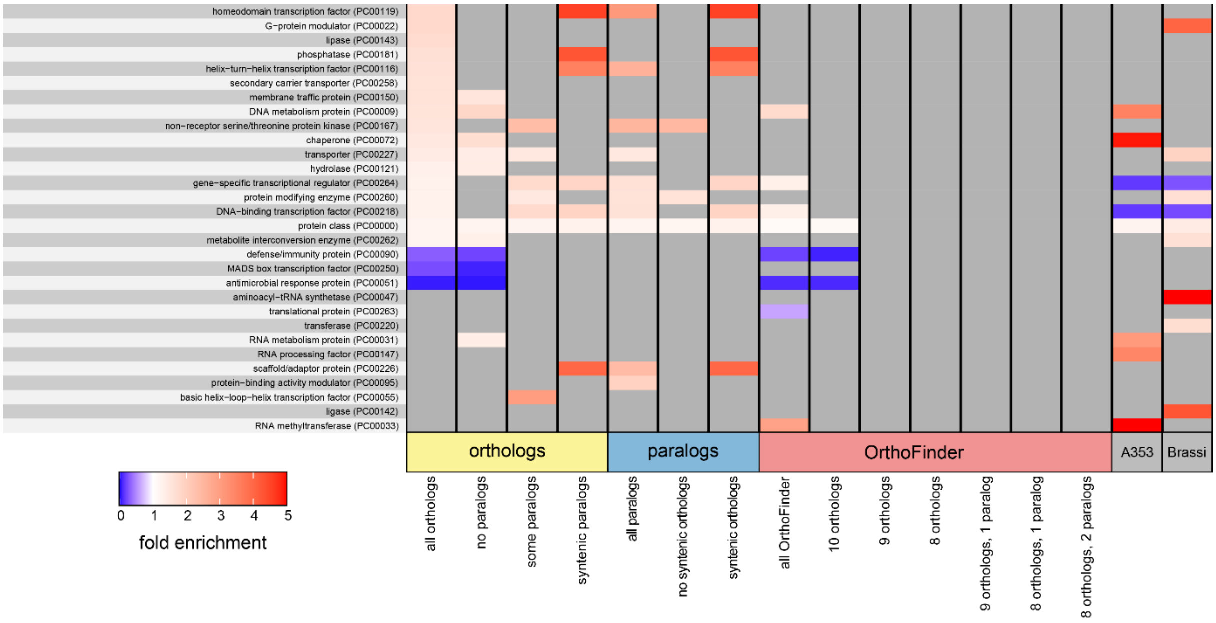


**Fig. S6. Enrichment for PANTHER protein class.** Fold enrichment for terms with significant enrichment (binomial test with Bonferroni correction for multiple testing; p-value < 0.05) are shown for all 16 gene sets. Heatmap was plotted using R package superheat v1.0.0 (Barter and Yu, 2022), fold enrichment > 5 was rare and thus merged.

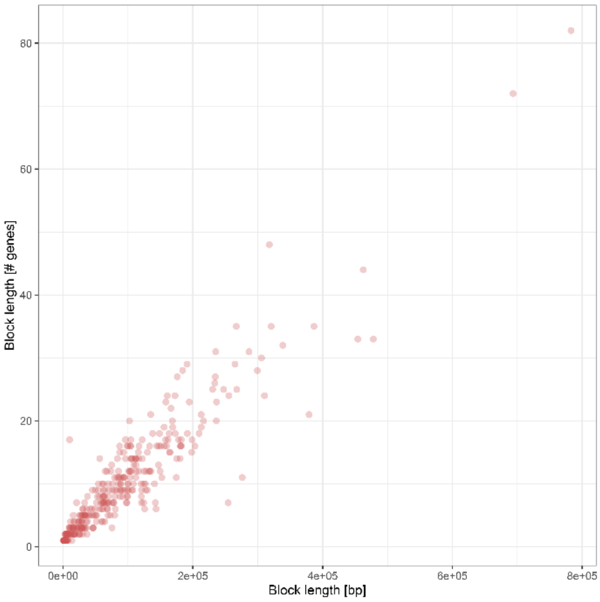


**Figure S7. Syntenic block length for ancestral genome reconstruction.** Block length in base pairs (bp) relative to number of genes for blocks used for ancestral genome reconstruction.

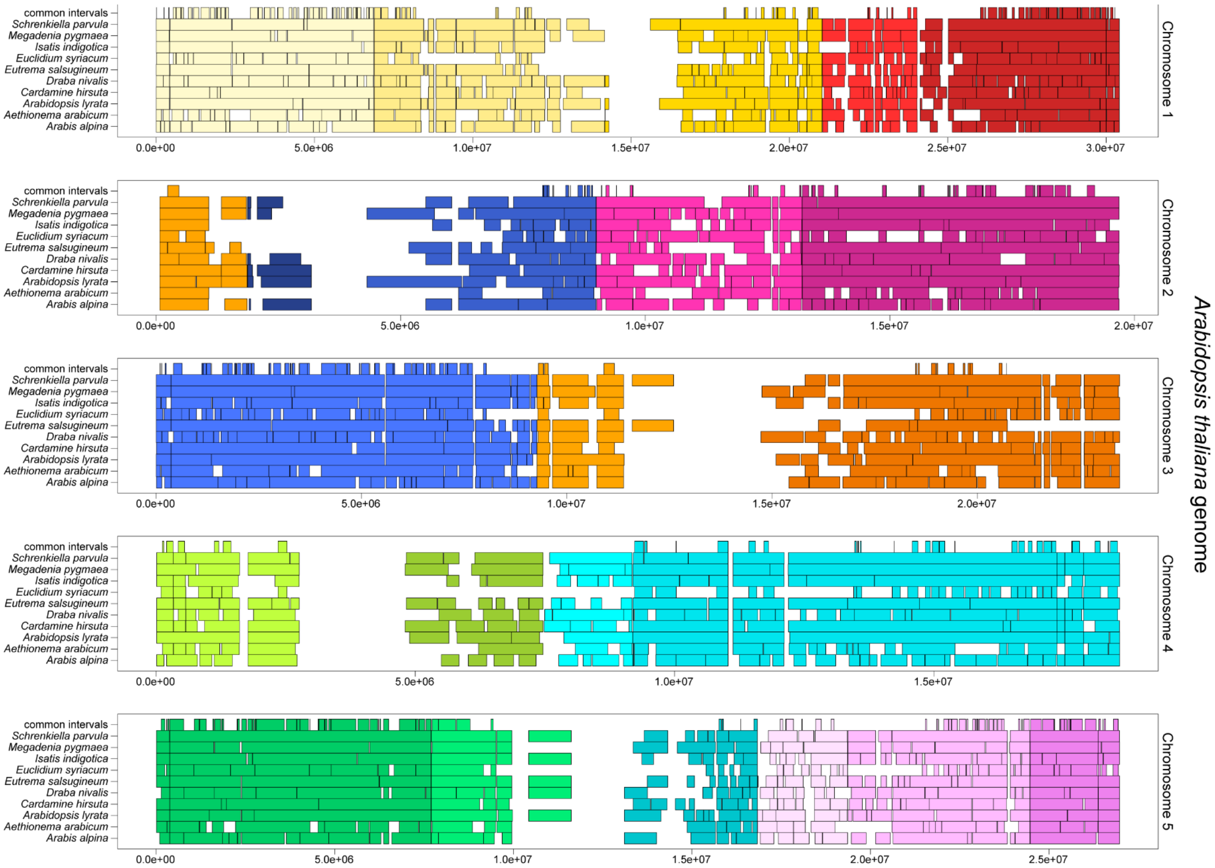


**Figure S8. Genomic coverage of syntenic blocks.** Pairwise syntenic blocks to the genome of *Arabidopsis thaliana* are displayed relative to the five *A. thaliana* chromosomes using the boundaries of blocks containing the genes common across Core Brassicaceae. Colors represent ABC blocks following Lysak et al (2016).

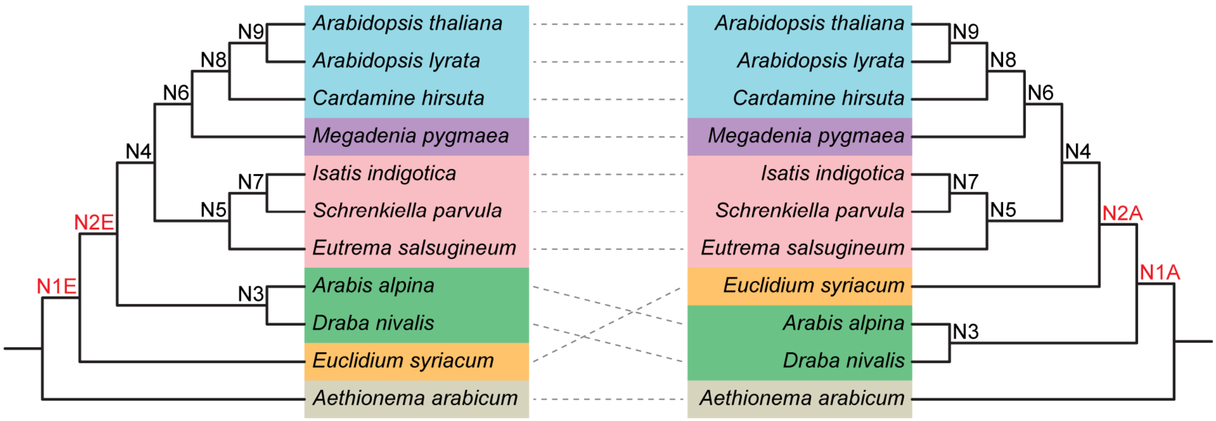


**Figure S9. Input phylogenies for ancestral genome reconstruction**. The position of clade D (*Arabis alpina*, *Draba nivalis*) and clade E (*Euclidium syriacum*) is different between both topologies, resulting in two different nodes (N2, N1).

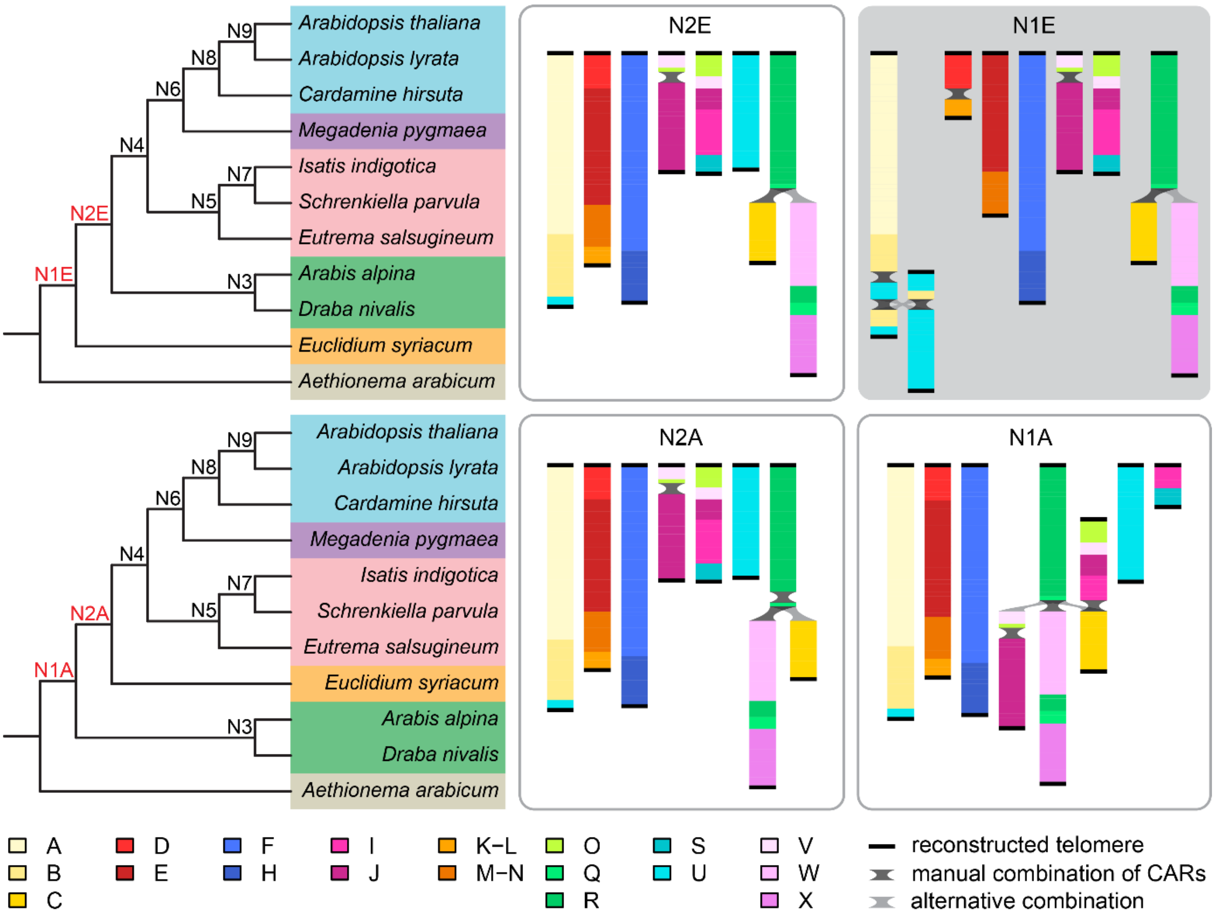


**Figure S10. Schematic illustration of ancestral genomes at nodes N1 and N2**. Markers are represented by horizontal bars colored by ABC blocks (Schranz et al., 2006; Lysak et al., 2016), black bars represent telomeres reconstructed by ANGES. CARs without two reconstructed telomeres were combined manually to retain marker adjacencies also found in related extant genomes; when multiple combinations were possible, the one found in the respective first diverging lineage (*Euclidium* or Arabideae) is displayed in dark grey, while the alternative combination is displayed in light grey. Reconstructions with *Euclidium* as first diverging lineage are shown in the top row and with Arabideae in the bottom row. The ancestral genome at N1E corresponds to the Core Brassicaceae ancestral genome (Fig. 5) and is highlighted in grey. Note that the displayed length of chromosomes here is relative to marker number, not nucleotide sequence or gene number.

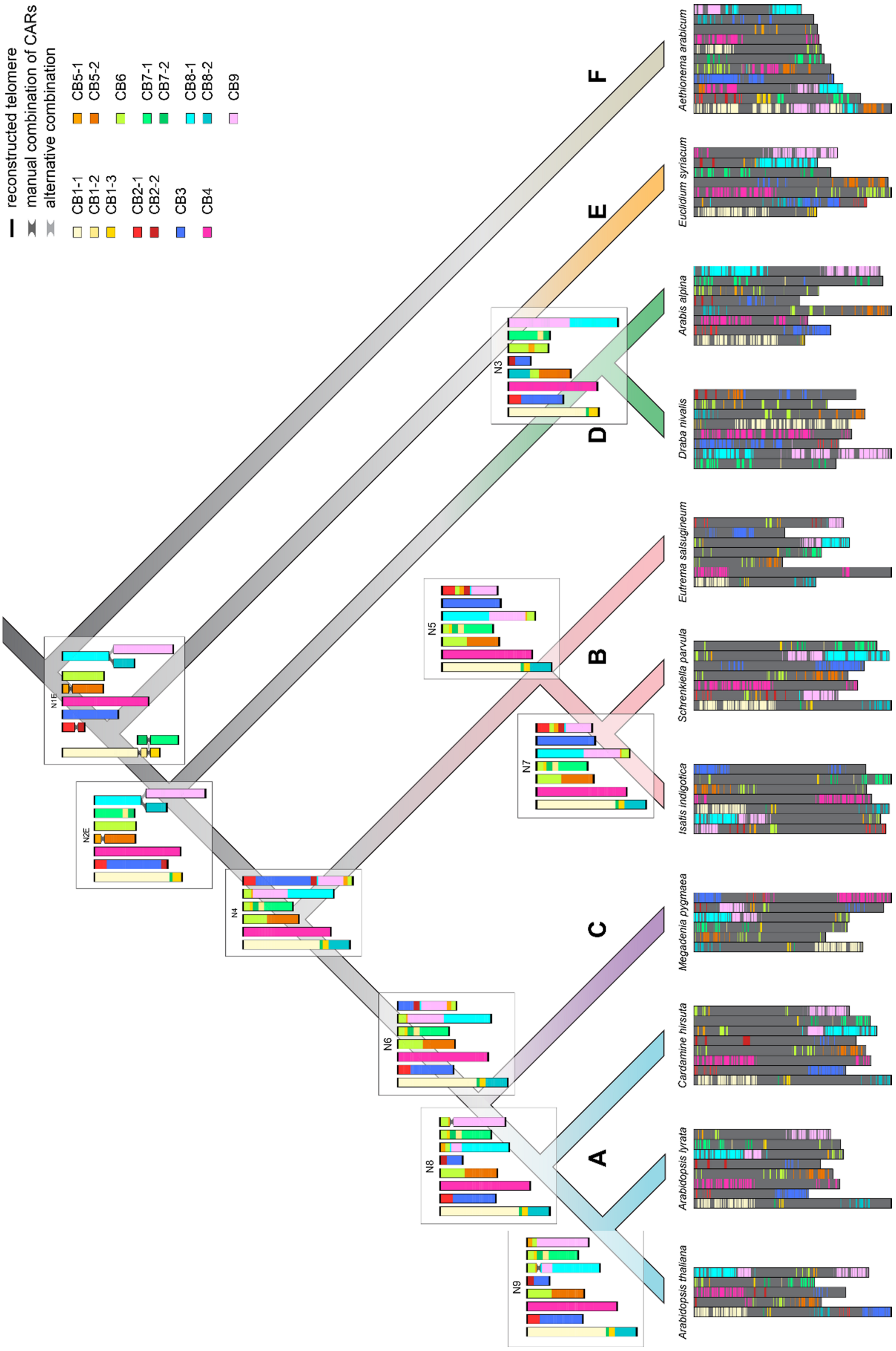


**Figure S11. Schematic illustrations of ancestral genomes along the Brassicaceae phylogeny.** Markers are represented by horizontal bars colored by CARs in the N1E reconstruction, black bars represent telomeres reconstructed by ANGES. CARs were combined manually for reconstructions containing CARs with less than two telomeres. Where more than one chromosome needed manual assembly, CARs were combined to retain marker adjacencies also found in related extant genomes; when multiple combinations were possible, the one found in the respective first diverging lineage is displayed in dark grey, while the alternative combination is displayed in light grey. Note that the displayed length of chromosomes here is relative to marker number, not nucleotide length or gene number except for the extant genomes.

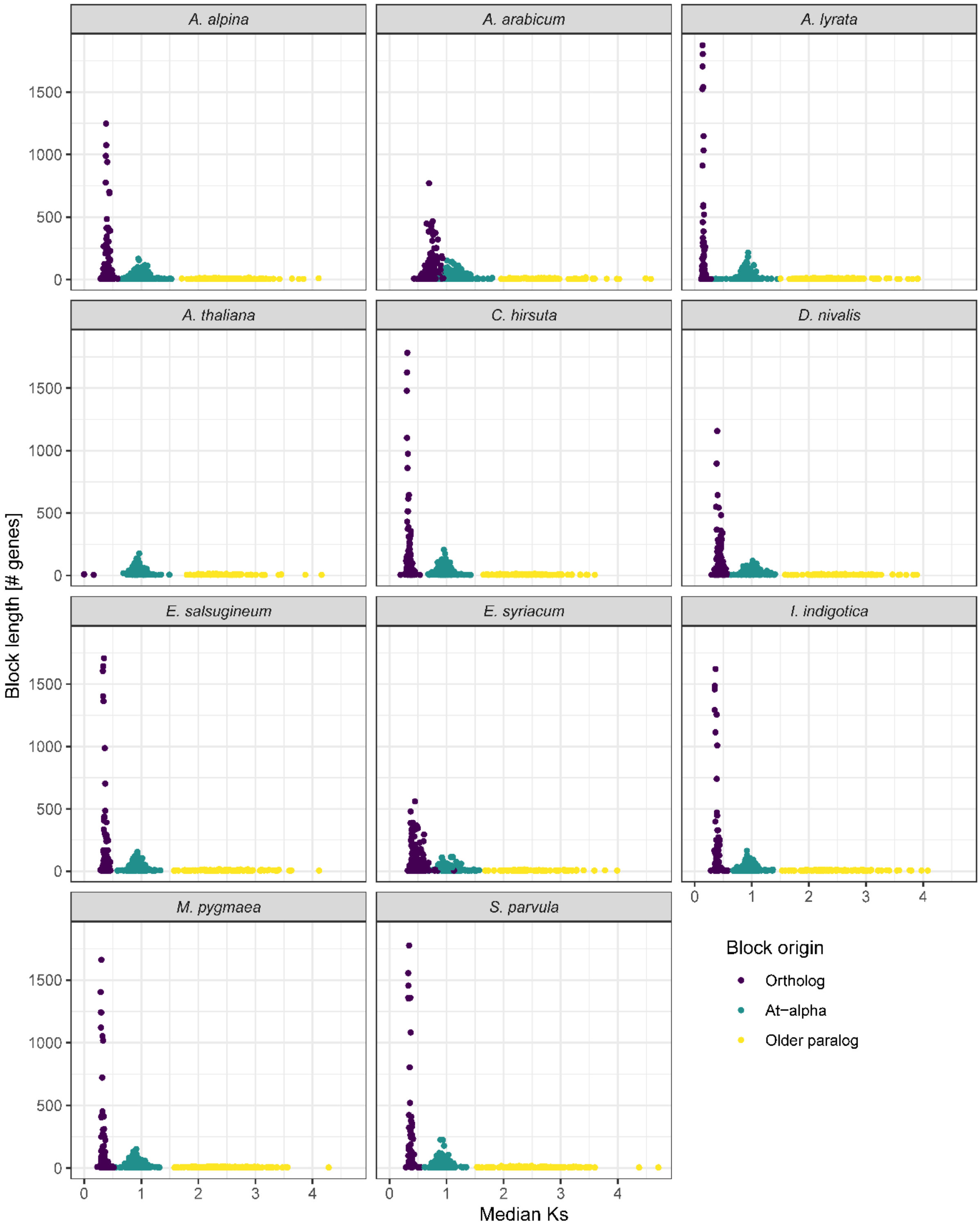


**Figure S12. Median block ks.** Median ks of syntenic blocks detected in CoGe SynMap relative to *Arabidopsis thaliana* are given for all ten other species. Median block ks was used as the threshold to determine orthology and paralogy of syntenic blocks using the species-specific thresholds indicated as a red dashed line. For *Aethionema arabicum* and *Euclidium syriaum*, block length and duplication status were also taken into consideration when assigning orthology/paralogy manually, as there were no clearly separated peaks for orthologs and At-α derived paralogs.
